# Molecular adaptation reflects taxon-specific mutational biases

**DOI:** 10.1101/2025.09.03.674101

**Authors:** Bryan L. Gitschlag, Arlin Stoltzfus, David M. McCandlish

## Abstract

A fundamental question in molecular evolution is the extent to which patterns of adaptive change are shaped by mutational biases that make some variants more likely than others to arise. Past studies provide support for important effects of mutation bias on adaptive change, but leave open the empirical question of how strongly and how broadly evolutionary patterns depend on taxon-specific mutational tendencies. To characterize this effect quantitatively, we aggregated nucleotide substitution spectra for adaptive amino acid changes from 14 species, primarily from adaptive laboratory evolution studies. We then paired each species-specific spectrum of adaptive changes with independently sourced measurements of the mutation spectrum of that same species. Across the 14 species, we find considerable heterogeneity in the relative rates of the six possible types of single-nucleotide changes, for both the mutational and adaptive spectra. Comparing these spectra across species, we find that, the stronger the bias toward a given mutation type in the mutation spectrum of a species, the more that mutation type contributes to adaptive substitutions in that species. We conclude that, for the adaptation of proteins via amino acid changes, taxon-specific patterns of adaptive substitution are strongly responsive to taxon-specific mutational biases across mutation types.

**Significance statement:** The rates at which different types of mutation occur vary widely, both for different types of mutations within a single species and across species. Historically, a strong concordance between patterns of sequence evolution and mutational biases was used to support the thesis that most sequence evolution is caused by variants that do not affect fitness. However, more recent evidence shows that mutational biases can also influence adaptive evolution. Using data from 14 species, we find a strong relationship between the rate at which different kinds of nucleotide mutations emerge and their rate of contribution to adaptive amino acid changes during protein evolution. These results suggest that processes that bias the production of variation have a strong impact even during adaptive evolution.

## Introduction

Broad patterns of gene and genome evolution reflect what is mutationally likely, an observation that has historically been taken as evidence for neutral evolution [1]. Yet population-genetic theory also suggests conditions under which such biases in mutation shape the course of adaptation, since the tendency of mutationally favored alleles to appear earlier than other alleles, i.e., arrival bias, makes them more likely to contribute to adaptation [2, 3]. The notion that mutational or other variational biases are a key factor in explaining evolutionary patterns, such as parallelisms and taxon-specific evolutionary tendencies, has persisted as a minority view for over a century [4, 5, 6, 3], despite a lack of robust evidence that such biases actually shape the course of adaptation. However, recent work has addressed this lack of evidence, establishing that, for a given species or taxonomic group, mutational biases often have a substantial effect on patterns of adaptive change [7, 8, 9, 10, 11, 12]. Nucleotide transitions and changes at CpG sites, for example, both occur at higher rates and contribute more to parallel adaptation [13, 14, 15, 16]. In another line of evidence, direct manipulation of the mutation spectrum during experimental evolution has a strong effect on which mutations serve as the drivers of adaptation, both when the spectrum is manipulated genome-wide [8] or altered locally by modulating the strength of a mutational hotspot [11] (for a review, see [17, 18] or Ch. 9 of [3]).

Importantly, the underlying theory for arrival bias predicts not merely an excess of likelier mutation types but systematically graduated effects. In other words, the greater the mutational bias, the greater the enrichment for that type of mutation amongst evolved adaptive changes, other things being equal [2, 17, 12]. Although prior empirical work by Cano *et al*. [12] comparing patterns of adaptation and mutation in three species suggests that the spectrum of adaptive changes within a species quantitatively reflects the mutation spectrum for that species, we do not currently know how broadly and how strongly patterns of adaptive substitutions respond (statistically) to species-specific mutational biases.

Here we address this question using data from 14 species for which published studies provide both (1) a nucleotide mutation spectrum and (2) sufficient data on adaptive changes to support a statistical analysis. We gathered empirical measurements of the mutation spectrum for each species, relying primarily on mutation-accumulation studies, together with independently sourced adaptive amino acid substitution data. Both the mutation spectra and the spectra of adaptive changes show substantial taxonomic heterogeneity; for example, transition-transversion bias varies approximately 10-fold across our 14 species. Across all species and datasets, we found a strong and systematic pattern of correlation: mutation types with higher rates contribute more to adaptation, and mutation types with lower rates contribute less. This pattern applies to the heterogeneity in rates across the six nucleotide mutation classes, the heterogeneity in the rate of individual mutation types across species, and heterogeneity across both species and mutation classes. This pattern remains robust and consistent under various challenges designed to uncover confounding effects of selection, non-adaptive contamination, and taxonomic sampling. We conclude that, for the adaptation of proteins via amino acid changes, taxon-specific patterns of adaptive substitution are strongly responsive to taxon-specific mutational biases across mutation types.

## Results

In this study, we sought to compare *de novo* mutation rates with patterns of adaptive change. To accomplish this, we assembled a novel, multi-taxon dataset that pairs a mutation spectrum with an independently sourced spectrum of adaptive substitutions, for each species for which such data were available in the published literature. Using a codon-based model of single-nucleotide mutation enabled us to specify a spectrum of relative adaptive substitution rates for each of the six types of single-nucleotide mutation, controlling for the codon composition of protein-coding regions in each species and the structure of the amino acid code (Fig. 1A). To ensure that the adaptive data consist of *bona fide* adaptive changes, with minimal contamination by non-adaptive mutations, we restricted the dataset to include only amino acid substitutions (excluding synonymous, noncoding, and nonsense changes) that meet multiple distinct criteria of evidence for having an adaptive effect (see Methods for details). In most cases, these data were obtained from experimental evolution studies, in which organisms were exposed to a selection pressure, such as a xenobiotic stressor or a novel niche, followed by genetic sequencing to identify mutations conferring the observed adaptive phenotype.

**Fig 1:**
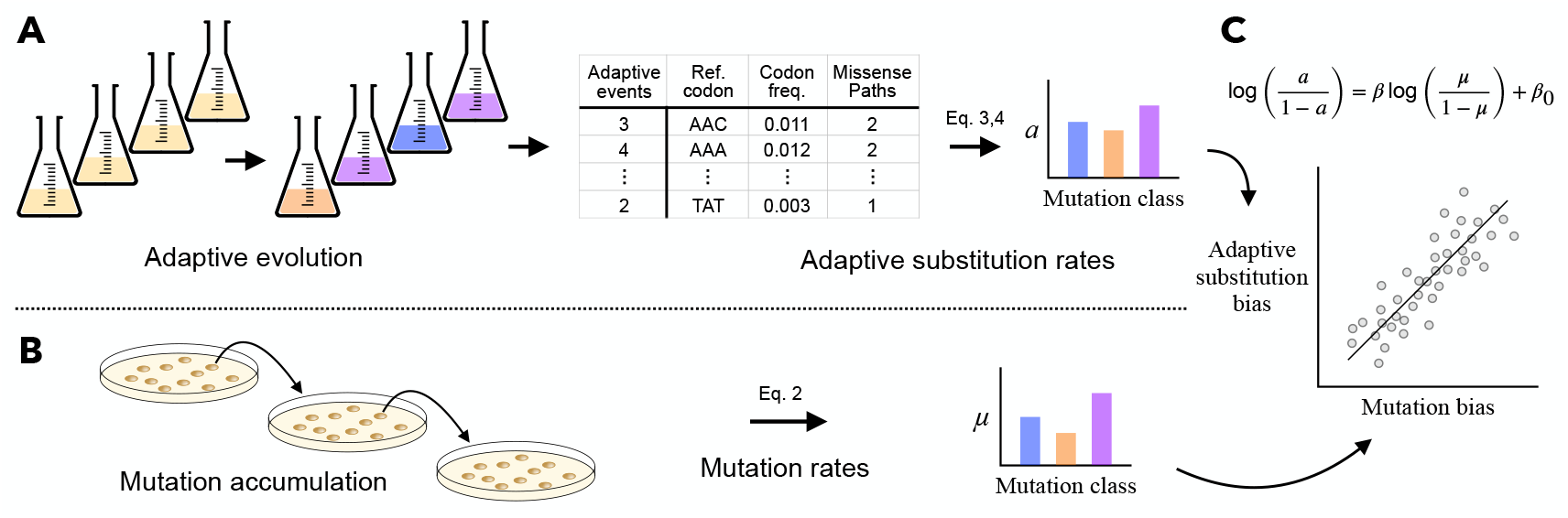
Schematic of workflow to assess relationship between molecular adaptation and mutation bias. A) Published reports of adaptive evolution traced to missense single-nucleotide substitutions (left) supplied observed counts for adaptive substitutions (*Adaptive events* column), which were transformed (using codon frequencies and accessible missense paths), into normalized adaptive substitution rates (*a* values at right; see Eq. 3, 4 in Methods for details). B) Published studies of mutation accumulation and segregating neutral variation (left) supplied counts of nucleotide mutations, which were transformed using base composition into normalized mutation rates (µ values at right). C) For a set of 14 species where both kinds of data were available, we evaluated the relationship between adaptive substitution bias and mutation bias via linear regression (Eq. 1).

Next, we curated a dataset of *de novo* mutation spectra, for each species for which the mutation spectrum could be paired with an adaptive substitution spectrum. For each species, the mutation spectrum represents the distribution of rates at which each of the six types of single-nucleotide mutation occurs, corrected for species-specific genome nucleotide composition. Mutation spectra are obtained from published mutation-accumulation experiments for most species (Fig. 1B), or from patterns of segregating neutral variation in the case of *Mycobacterium tuberculosis* and *Toxoplasma gondii*, and exhibit considerable taxonomic heterogeneity, consistent with prior studies [19].

Combining the data on adaptive substitutions with the data on *de novo* mutations resulted in a dataset covering 14 species, which includes, for each species, a spectrum of adaptive substitution rates based on observed adaptive amino acid changes (median 36 adaptive events) and a nucleotide mutation spectrum obtained from independent sources (median 245 *de novo* mutations, Table 1 and Fig. 2; see Methods). Consistent with Cano *et al*. [12], comparing the two types of spectra reveals significantly positive intraspecific correlations between mutational bias and bias in adaptive substitutions for 13 of the 14 species (Fig. S1 and Table S1), suggesting that within any given species mutational classes with higher rates tend to also contribute to adaptation at higher rates. Thus, with the enlarged multi-species dataset used here, we extend the finding of Cano *et al*. [12] to roughly four-fold more species.

**Table 1:** Sample sizes for mutational and adaptive events for each species in this study. Sizes of original datasets used in this study, consisting of the number of observed adaptive events from which the spectra of adaptive substitutions are derived (Fig. 1A), together with the number of observed mutation counts from which the *de novo* mutation spectra are derived (Fig. 1B).

| Species | *De novo* mutations |  | Adaptive substitutions |  |
| --- | --- | --- | --- | --- |
|  | Counts | References | Counts | References |
| *B. cenocepacia* | 245 | [20] | 36 | [21, 22, 23] |
| *B. subtilis* | 350 | [24] | 29 | [25, 26, 27] |
| *C. glutamicum* | 240 | [28] | 22 | [29, 30, 31, 32] |
| *D. discoideum* | 37 | [33] | 22 | [34] |
| *E. coli* | 312 | [35, 36] | 301 | [37, 38, 39, 40] |
| *L. lactis* | 813 | [41] | 28 | [42, 43, 44, 45] |
| *M. tuberculosis* | 104,622 | [12] | 4,414 | [12] |
| *P. aeruginosa* | 44 | [46] | 144 | [47] |
| *P. falciparum* | 117 | [48] | 84 | [49] |
| *P. fluorescens* | 253 | [50] | 35 | [51, 52] |
| *S. aureus* | 274 | [41] | 104 | [53] |
| *S. cerevisiae* | 1,987 | [54, 55, 56] | 202 | [57, 58] |
| *S. enterica* | 80 | [59] | 30 | [60] |
| *T. gondii* | 97 | [61] | 37 | [62, 63] |

**Fig 2:**
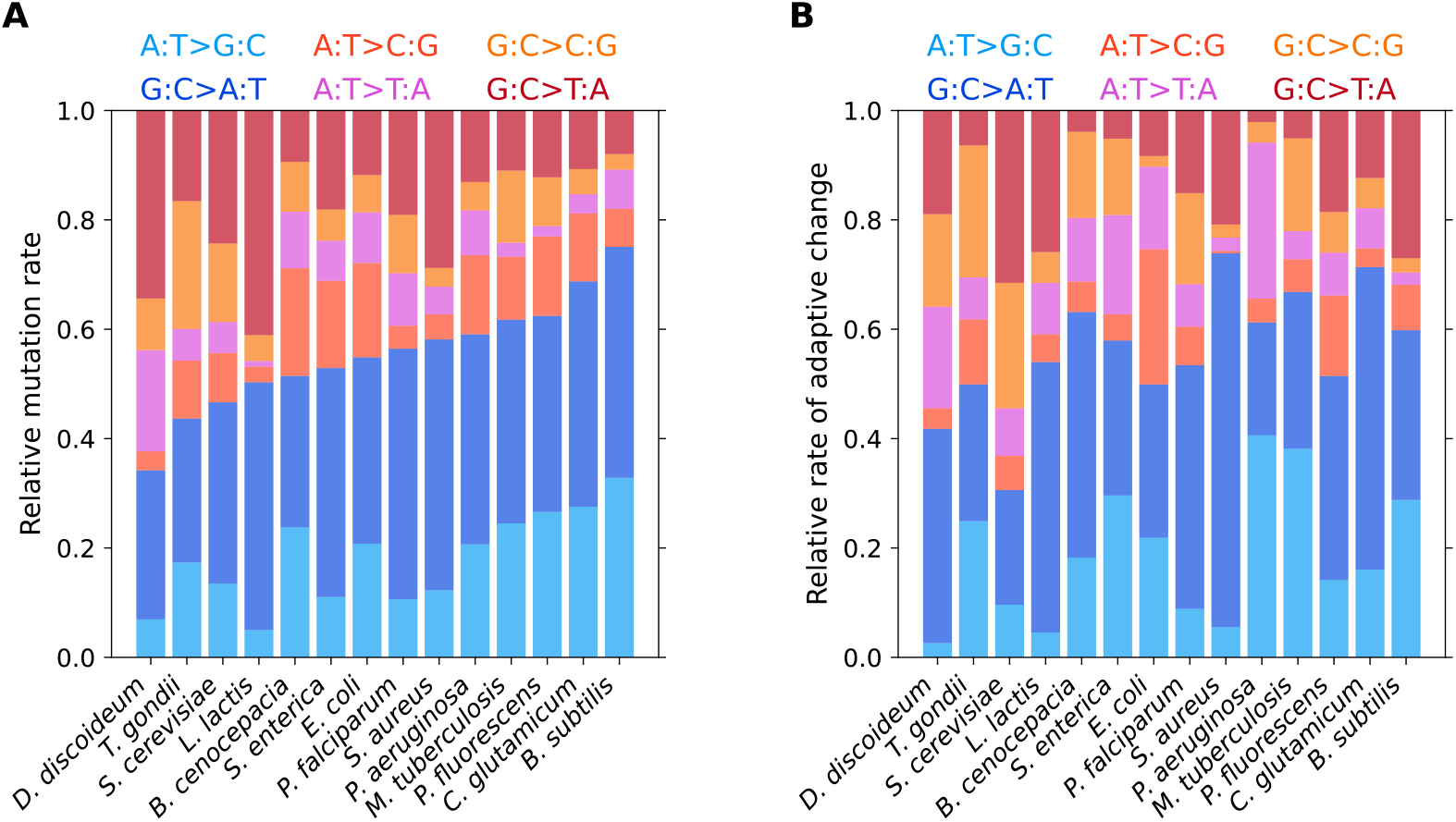
Graphical summary of mutation spectra and spectra of adaptive substitutions by species. A) Spectra of relative de *novo* mutation rates for 14 species, arranged in order of increasing ratio of transitions (cool colors) to transversions (warm colors). B) Spectra of adaptive substitution rates by species, arranged to preserve the ordering of species in panel A.

### Transition bias of adaptive substitutions reflects species-specific *de novo* transition bias

A distinct issue from the one addressed in Cano *et al*. [12], and our main focus here, is how interspecific differences in mutation bias affect interspecific differences in adaptive evolution. In other words, when we look across species, do species with particularly strong mutational biases also exhibit strongly biased patterns of adaptive substitution? To address this question, we begin by considering transition bias, defined as the ratio of the rate of transitions (purine-to-purine or pyrimidine-to-pyrimidine) to transversions (purine-to-pyrimidine or vice versa). Although transitions tend to be overrepresented among adaptive substitutions [64, 65], the cause for this remains a topic of debate, with some studies attributing transition-biased adaptation to selection against transversions [66]. To more systematically investigate the effect of biased mutation rates, we fit a linear regression by least squares on the log-transformed data,

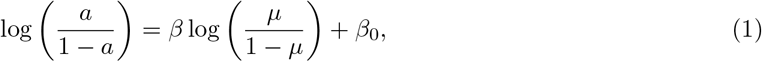

where µ and *a* are the sums of the two transition components of the normalized mutation and adaptive substitution spectra, respectively, so that µ/(1 − µ) and a/(1 − a) are the ratios of the summed transition to transversion rates in the corresponding spectra, *β*_0_ is the intercept, and *β* is a statistic that captures the effect of mutation bias on adaptive substitutions, such that *β* = 1 indicates a directly proportional relationship between mutation bias and adaptive substitution bias. Thus, the question that we are asking is whether the ratio of summed transition to transversion rates in the mutation spectrum of a given species predicts the corresponding ratio in its spectrum of adaptive amino acid substitutions. We observe a strong positive effect of mutation bias (*β*=0.67, 95% bootstrap confidence interval 0.28 to 1.11; Fig. 3, gray lines show bootstrap regressions), and this effect of transition bias on adaptive change is significant when compared against the regression on simulated data under the null model in which transition bias has no effect on adaptive change (p=0.0012; see also Table S2). We conclude that transitions tend to contribute more to adaptive change in the species in which transitions occur at a higher rate.

**Fig 3:**
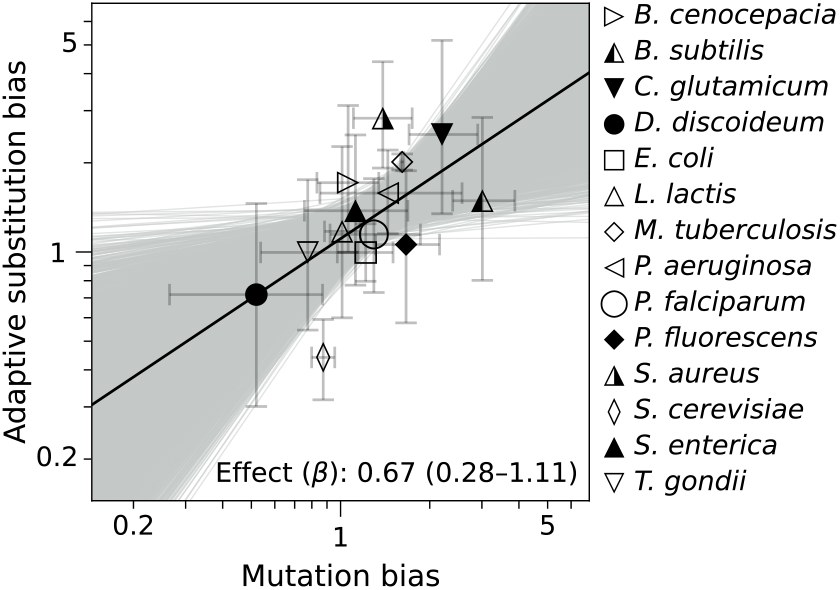
Transitions contribute more toward adaptive changes in species where transitions occur at a higher *de novo* rate. For each of the 14 species in Table 1 and Fig. 2, the ratio of rates of transitions to transversions among adaptive amino acid substitutions is plotted against the ratio of rates of *de novo* transitions to transversions. Least squares linear regression on log-transformed empirical data (solid black line) and 10,000 bootstrap samples (gray lines). The 95% bootstrap confidence interval for mutation bias effect, *β*, is shown in parentheses. Significance of *β* was assessed by simulating datasets (n=10,000) under the null model in which mutation bias has no effect on bias in adaptive substitutions, p=0.0012.

### Mutation bias predicts bias in adaptive outcomes across taxa and mutational classes

If our results so far represent a generalizable effect of mutation bias, rather than a peculiar feature of transitions, then we expect a similar relationship between mutation bias and adaptive substitution bias regardless of mutation type. To explore this possibility, we re-expressed the mutational data as six separate biases for each species (one bias per class of possible single-nucleotide mutation), so that if a mutation class occurs at rate µ in the mutation spectrum and at rate *a* in the adaptive spectrum then we calculate the corresponding biases as µ/(1 − *µ)* and a/(1 − *a*). Pooling data over all mutational classes and all species and estimating the coefficient *β* using Eq. 1 as before, we again found a strong positive relationship between mutation bias and adaptive substitution bias (*β* =0.8, 95% bootstrap confidence interval 0.73 to 0.91, p<0.0001 under a null model where biases in mutation rate have no effect on adaptive substitutions; Fig. 4, Table S2).

**Fig 4:**
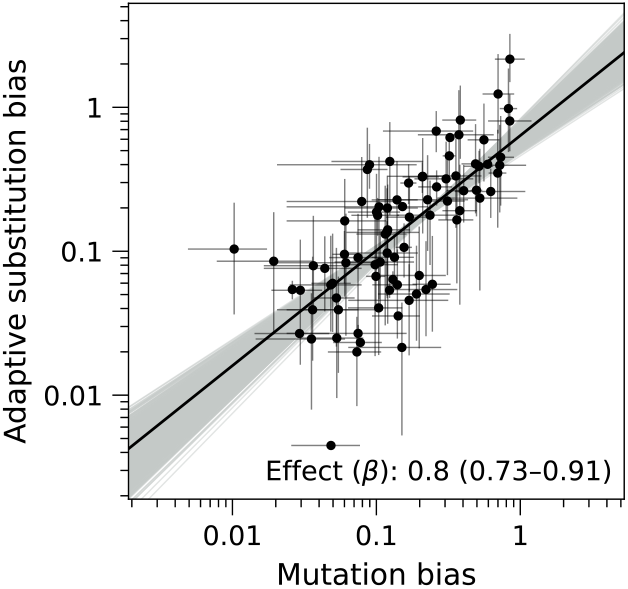
Mutations that occur at a higher *de novo* rate contribute more toward the rate of adaptive substitutions when aggregating data across taxa and mutational classes. Data points (n=84) represent each of the six single-nucleotide mutation classes in each of the 14 species from Table 1 and Fig. 2. Values of mutation bias and adaptive substitution bias are expressed as rate ratios for each mutation class and species. Least squares linear regression on log-transformed empirical data (solid black line) and 10,000 bootstrap samples (gray lines). The 95% bootstrap confidence interval for the mutation bias effect, *β*, is shown in parentheses. Significance of *β* was assessed by simulating datasets (n=10,000) under the null model in which mutation bias has no effect on bias in the rates of adaptive substitutions, p<0.0001.

### Isolating interspecific variation in bias for distinct mutation classes

The analysis in Fig. 4 combines intra- and interspecies variation in mutational and adaptive spectra into a single regression. In order to conduct an analysis where our coefficient *β* captures only the interspecific component of the signal, we fit two additional models. These models both have separate intercepts for each mutational class, where these class-specific intercepts capture the taxon-independent bias for or against each mutational class among adaptive substitutions. More concretely, we note that some mutational classes have consistently higher mutation rates than others across all species (e.g. G:C>A:T generally occurs at a high rate and A:T>T:A occurs at a low rate regardless of species). Thus, our goal here is to determine whether the correlation observed in Fig. 4 simply reflects enrichment of generally high-rate mutation classes among adaptive substitutions, or whether mutational biases also predict adaptive biases across species within each mutation class. For example, do species with particularly strong mutational biases towards G:C>A:T also exhibit particularly strong biases towards G:C>A:T among adaptive substitutions?

In the first model, all six mutation classes share a single *β* parameter but have separate class-specific intercepts *β*_0_, (see Methods, Eq. |5), so that *β* captures the impact of interspecific variation in mutation spectrum on the spectrum of adaptive substitutions, while the *β*_o,*i*_ capture factors that influence the adaptive spectrum in a species-independent manner. Under this model, we estimate that *β* = 0.74 (95% bootstrap confidence interval from 0.6 to 0.9, p<0.0001, Fig. |S2), indicating a strong and almost proportional influence of interspecific variation in mutational spectrum on the spectrum of adaptive substitutions.

In the second model, we fit a separate regression model for each mutation class, allowing the effect of mutation bias on adaptive substitutions to vary for each type of mutation (that is, Eq. 1 is fit to the data for each mutation class individually, producing a separate *β* value for each class). All 6 regression models had significantly positive slopes relative to a null model where mutation rates have no impact on the spectrum of adaptive substitutions (Fig. 5, Fig. |S3, and Table S2, *p* ≤ 0.022 for all slopes against simulated data under the null model in which mutation bias has no effect). More broadly, the strong degree of overlap between the different regression lines in Fig. 5 suggests a qualitatively similar relationship, regardless of mutation type, between the degree of mutation bias and the degree of bias in the spectrum of adaptive substitutions, consistent with the hypothesis that mutational biases play a strong role in shaping the spectrum of adaptive substitutions.

**Fig 5:**
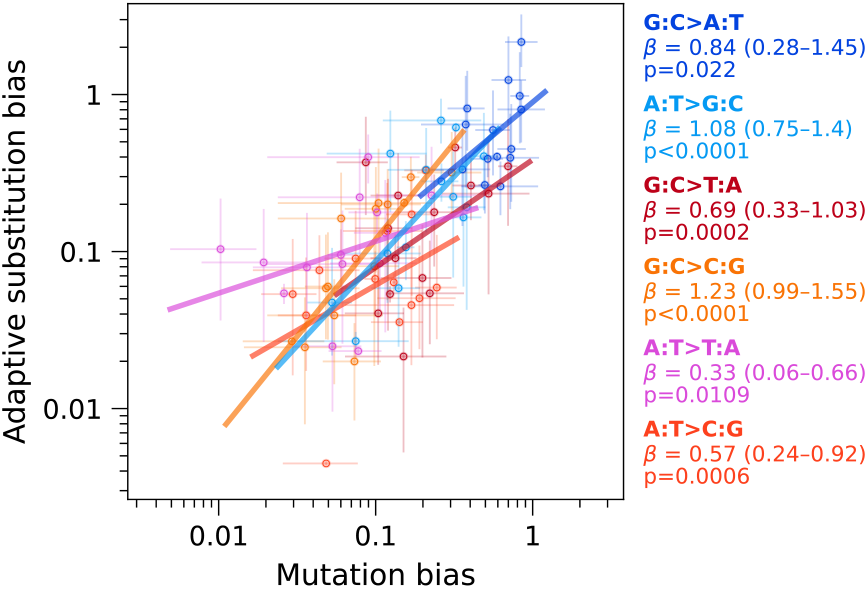
Variation across species in adaptive substitution bias reflects variation in mutation bias for each of the single-nucleotide mutation classes. Bias in the rates of adaptive substitutions is plotted against bias in mutation rates, across 14 species, for each of the six mutation classes individually, via least squares regression on the log-transformed data (solid lines). Error bars show 95% bootstrap confidence intervals (n=10,000 bootstraps) for mutation bias and adaptive substitution bias. The 95% bootstrap confidence interval for mutation bias effect, *β*, is shown in parentheses for each mutation class individually, together with p-values for *β* assessed by simulating datasets (n=10,000) under the null model in which mutation bias has no effect on bias in adaptive substitutions.

### Mutation bias predicts both sites of adaptive changes and the identity of the mutant base

Biases in the rates of different types of mutations can be influenced by two distinct aspects of the mutation process. Some nucleotide bases may tend to be more mutationally volatile than others, a phenomenon we call site bias. At the same time, mutations starting from a given base may tend to prefer certain alternate bases, which we refer to as alternate-base bias. Does the spectrum of adaptive substitutions reflect site bias, alternate-base bias, or both?

To address this question, we repeated the regression analyses in a manner that partitions the mutation spectra and adaptive spectra into site bias and alternate-base bias. Under our six-rate spectra, site bias corresponds to whether G:C or A:T sites are more likely to be changed (see Methods). We observe a roughly four-fold range of mutation site biases for most species, with the exception of *L. lactis*, which had a ten-fold bias in favor of mutations occurring at G:C sites. Nonetheless, we observe a significant positive effect of site bias on adaptive outcome (*β*=0.83, 95% bootstrap confidence interval 0.57 to 1.11, p<0.0001 against a null model in which mutation bias has no effect on adaptive substitutions; Fig. S4). Likewise, alternate-base bias in mutation varies by approximately ten-fold between species, and we observe a strong positive effect on adaptive outcome (*β*=0.7, 95% bootstrap confidence interval 0.51 to 0.9, p<0.0001 against a six-intercept null model using Eq. 7; Fig. S5). Thus, both site bias and alternate-base bias in mutation influence the adaptive spectrum.

### Assessing the effect of taxonomic sampling

We sought to follow up on our findings by assessing their robustness to potential sources of error. From the publicly available data, we were able to pair a *de novo* mutation spectrum with a spectrum of adaptive changes for 14 species. Due to the relatively small set of species, we considered the possibility that a single outlier species might be responsible for the apparent effect of mutation bias on outcomes of adaptive evolution.

How might such an outlier affect our analysis? To address this question, we repeated the previously described regression analyses while omitting each of the 14 species one at a time. First, we conducted a leave-one-species-out cross-validation analysis. Specifically, we asked whether fitting the model in Eq. 1 to 13 of the 14 species could be used to predict the adaptive substitution biases for the held-out species from its observed mutation biases. Fig. 6A aggregates these predictions over all 14 species, showing a Pearson correlation of 0.68 between our predictions and the observed biases among adaptive substitutions. Conducting a species permutation test of the empirical adaptive substitution spectra, we found that predicted adaptive substitution biases correlate more strongly with the observed biases than they do with the biases under species-permuted adaptive spectra (p=0.0005). These results indicate that the relationship between mutational biases and biases among adaptive substitutions is sufficiently consistent between species to allow accurate prediction of the adaptive spectrum for held-out species.

**Fig 6:**
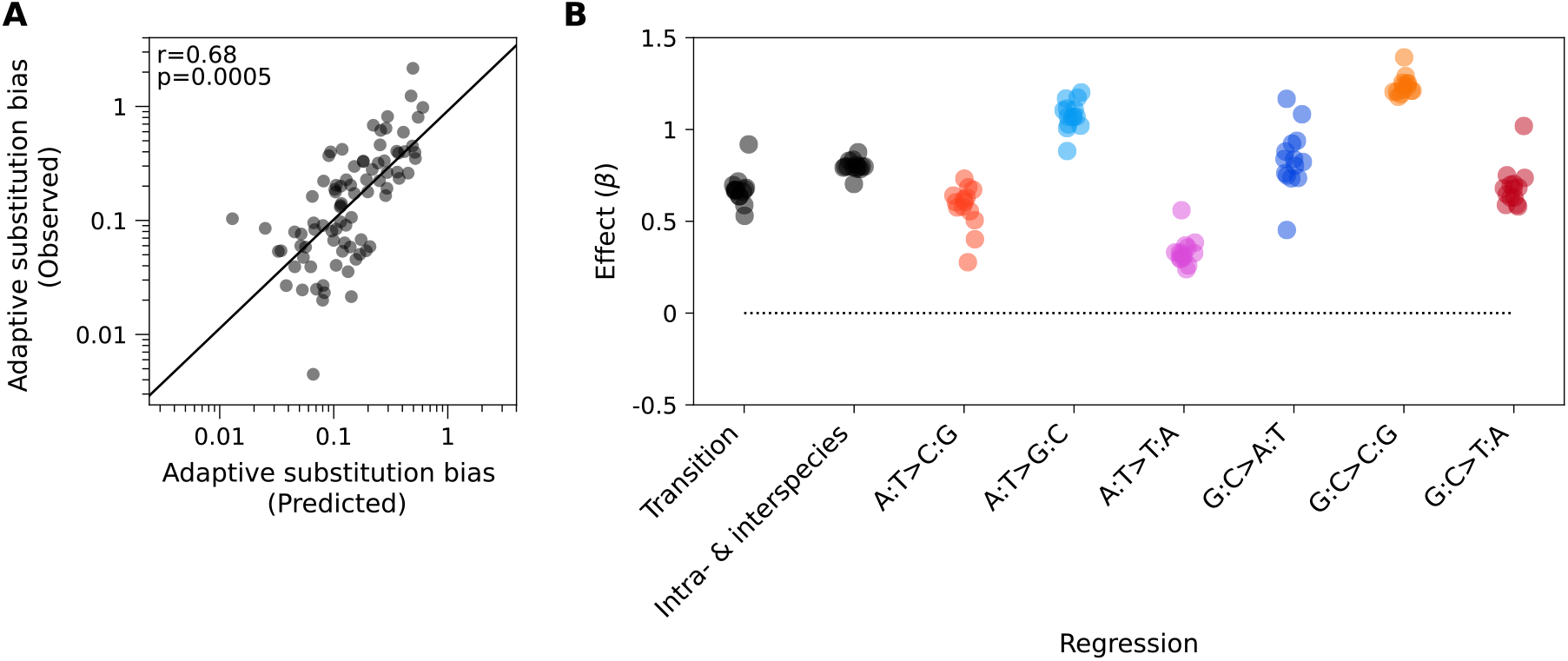
The relationship between mutation bias and adaptive substitution bias is robust under species exclusion. A) Observed versus predicted bias in rates of adaptive substitution under leave-one-species-out cross-validation, aggregated across all 14 species and all six mutation classes. For each held-out species, the predicted adaptive substitution bias was obtained using Eq. 1 fit to the 13 other species and applied to the empirically observed mutational bias from the held-out species. The p-value for the leave-one-species-out correlation was assessed by permuting the species labels of the observed adaptive spectra and computing the Pearson’s *r* between the log-transformed predicted and observed biases (n=10,000). B) Leave-one-species-out estimates of *β* obtained by repeating the regression analyses reported in (from left to right) Fig. 3, Fig. 4, and the individual regressions from Fig. 5, while omitting each of the 14 species one at a time. See also Table

Second, we examined the numerical estimates of *β* calculated when one species is held out. We found that the regressions for transition bias (Fig. 6B), overall intra- and interspecies variation in bias (Fig. 6B), bias for each of the six mutation classes (Fig. 6B), and interspecies variation in bias (Table S3), all produced positive estimates of *β*, regardless of which species was held out. These results indicate that our main conclusions are not driven by the data from any single species. With respect to the potential influence of phylogenetic non-independence in our dataset, the only moderately close species pairs are (1) *E. coli* and *S. enterica* with an average genomic nucleotide identity (ANI) of 80.9% and a divergence time on the order of 120 MYA [67], and (2) *P. aeruginosa* and *P. fluorescens* with an ANI of 75.8% and an unknown divergence time (ANI was calculated with OrthoANI Tool [68] using NC 000913.3 for *E. coli*, NC 003197.2 for *S. enterica*, NC 002516.2 for *P. aeruginosa* PA01, and NZ OV986001.1 for *P. fluorescens* SBW25). To remove any non-independence due to these pairs, we repeated our analysis using datasets missing one member of each pair, considering all 4 such combinations, finding that our estimates of *β* were robust to these exclusions (Fig. S6, Table S4).

### Accounting for taxon-specific uncertainties

Our regression analyses have so far been conducted using least squares, based on the assumption that deviations from the model fit are largely driven by species-specific biological factors. However, some of the underlying datasets in our analysis are small, suggesting that the residuals in our regression models could be driven by sampling noise in estimating the underlying rates. To explore this possibility, we used a weighted least squares (WLS) regression model (see Methods), which accounts for species-specific uncertainties in adaptive substitution bias by inversely weighting the residuals for each datapoint by the variance in the estimated log adaptive substitution bias. In order to account for uncertainty in the mutational biases, we also performed a weighted total least squares (WTLS) regression, which modifies the WLS approach to weight the residuals in both the *x* and *y* directions based on the variances for the estimated log *de novo* mutation biases and log adaptive substitution biases, respectively. Both the WLS and WTLS regressions show a highly significant positive association between adaptive substitution bias and mutation bias (p<0.0001), for each of transition bias, overall bias across taxa and mutational classes, and the six individual mutational classes (Fig. S7, S8). We conclude that our original findings are robust to the stochastic effects that result from having small sample sizes for some of the species in our dataset.

### Assessing the effect of non-adaptive contamination

We also considered whether contamination of the adaptive spectra by non-adaptive changes could impact our conclusions. Although we applied stringent filtering criteria to remove mutations for which we do not have substantial evidence of an adaptive effect (see Methods), the resulting dataset may still contain some neutral or deleterious hitchhikers. In the case of thermal adaptation in *E. coli* [39], for example, if we were to remove the criterion that all included changes must be missense mutations then four synonymous changes would have met the same criteria that were used to identify the 282 adaptive missense changes included in our study (see SI Methods). If we assume the synonymous changes to be neutral hitchhikers, their presence at 1.4% frequency (4 of 286) implies that approximately 3.6% of the missense changes are also neutral hitchhikers, given the relative rates of mutation adjusted for codon usage in *E. coli*. In contrast, zero synonymous changes would have met the same criteria that were used to identify the 202 adaptive missense changes in *S. cerevisiae*, suggesting an even lower level of contamination; likewise, in *M. tuberculosis*, all changes included in our dataset are known to confer antibiotic resistance [64].

In order to assess the robustness of our conclusions to even very high levels of non-adaptive contamination, we considered how levels of contamination of 10%, 20%, 30% and 40% would affect our results. For each of these possible levels of contamination, we removed the corresponding percentage of mutations from the adaptive dataset before repeating our analyses. These contaminants were assumed to be drawn from the mutation spectrum, hence the data removed from each mutation class was proportional to the frequency of that class among *de novo* mutations (see Methods), representing a worst-case scenario in which contamination contributes maximally to the apparent effect of mutation bias on adaptive outcome. For the analyses that jointly considered data from all six mutation classes, our estimates of *β* were highly consistent and remained significant relative to the null model even under the pessimistic assumption that 40% of the dataset consists of neutral hitchhikers (Fig. S9, top three rows). Although the impact of contamination was greater upon estimating *β* for individual mutation classes, *β* remained significant in the case of 20% contamination for all six mutation classes, and *β* remained significant in the case of 40% contamination for most mutation classes, including A:T>G:C and all classes of mutations at G:C sites (Fig. S9, bottom row). Given the stringency of our data curation and the robustness of our analyses to even implausibly high levels of contamination, we conclude that hitchhiker mutations are unlikely to explain the observed relationship between mutation bias and bias in adaptive outcome.

### Potential selective influences on the spectrum of adaptive substitutions

Though our analysis is motivated by a specific theory about the role of mutation in evolution, our approach is correlational, and thus it is reasonable to consider how much of the correlation could be explained by other influences, particularly selection (e.g., see Lyons and Lauring, [66]). For instance, our procedure for estimating the spectrum of adaptive substitutions treats all amino acid changes as *a priori* equally likely to contribute to adaptation whereas in fact there are systematic differences in exchangeability between amino acids [69, 70]. To incorporate the effect of such differences, we modified our regression analyses to include differential acceptance of amino acid changes based on Tang’s U [69], an empirical measure (based on protein divergence patterns) that is designed to capture differences in amino acid exchangeability while controlling for the effects of mutation and the genetic code. Including Tang’s U had a negligible effect on the regressions (Fig. S10), suggesting that our results are not driven by a failure to account for systematic differences in amino acid exchangeability. Another approach to addressing the influence of selection is to note that our intercept parameters *β*_0_ reflect the overall enrichment of any given class of mutations amongst adaptive substitutions relative to random mutations. While our null hypothesis so far has been a point-null based on the distribution of mutationally accessible missense changes in each species, we can also consider a stronger null that selection enriches or depletes for each type of mutation similar to what is observed in the data, but that *β* = 0 so that there is no linear relationship between log adaptive bias and log mutational bias. We conducted a one-sided test of this hypothesis by inverting our percentile bootstrap confidence intervals for *β*, finding that *β* is still significantly greater than 0 for all of our main analyses (Table S5, *p* ≤ 0.0073 for all fit values of *µ*).

A different selective hypothesis, suggested by [71], is that the observed alignment between mutation and adaptation reflects a history of selection shaping the mutation spectrum. For example under this hypothesis, when A to G mutations occur *de novo* at relatively higher rates in a species, this is because A to G mutations tend to be more beneficial in that species. One prediction of this hypothesis is that we would expect the mutation-evolution correlation to be weaker among neutral changes, as this correlation would reflect only mutational bias without the hypothesized selective effect. Therefore we examined data from 3 species for which our original datasets include synonymous changes, on the grounds that they are more likely to be neutral. The results shown in Fig. S11 indicate that, contrary to this alternative hypothesis, the mutation-evolution correlation for synonymous changes is, if anything, stronger (larger inferred *β*).

A final concern is that our results may reflect some artifact of combining data from heterogeneous selective conditions. Our leave-one-species-out analysis in Fig. 6 shows that our main results do not depend on conditions specific to any one species. To probe the effect of heterogeneity within selective conditions for a given species, we focused on the dataset from *M. tuberculosis*, which aggregates data across different antibiotics and target loci. Eight such subsets of the dataset are large enough to consider individually. This set of 8 includes one dataset not used in our main analysis because it consists of ribosomal RNA changes rather than amino acid changes. We find that adaptive substitution bias is positively correlated with mutation bias for each of these 8 datasets (Fig. S12). Thus, the observed mutation-adaptation correlation is robust to considerable heterogeneity in conditions of adaptation.

### Analyses using frequencies instead of rates

The associations we have reported so far are all based on normalized rates rather than raw frequencies. In particular, our rate calculation corrects for variation in the number of nonsynonymous mutational paths available for adaptation arising due to species-specific codon usage. However, in species with highly biased codon usage, substitutions that occur at high rates may nonetheless be infrequent among adaptive substitutions, e.g. A:T>G:C substitutions may occur at high rates but low frequencies in GC biased genomes because wild-type AT nucleotides are relatively rare. This raises the question of the relationship between mutational biases and biases in adaptive outcomes when measured on a frequency rather than rate scale.

To assess the effect of mutational biases on biases in the frequencies of adaptive outcomes, we repeated our previous regression analyses, with two important changes. First, raw adaptive substitution counts were converted directly to frequencies, rather than being corrected for codon usage. Second, mutation rates are also converted to predicted relative frequencies. With this approach, we find that biases in frequencies among adaptive outcomes and *de novo* mutations (Fig. S13) are significantly correlated with respect to intraspecific variation among mutational classes (Fig. S14 and Table S6), for the same 13 out of 14 species as observed in the case of biased rates (Fig. S1 and Table S1). We further note a significant positive association with respect to variation across species for transitions and the individual mutational classes (Figs. S15 to S17 and Table S7), as well as with respect to site bias and alternate-base bias (Fig. S18, S19). Similar to our analyses with respect to adaptive substitution rates, the association between mutation bias and bias in frequencies of adaptive outcomes is robust to the effects of taxonomic outliers in nearly all cases, with the exception of A:T>C:G upon omission of *E. coli* (Fig. S20 and Table S8). Moreover, our findings with respect to frequencies of adaptive outcome remain significant in the case of 10% contamination for all six mutation classes, 20% for A:T>T:A, 30% for G:C>T:A, and 40% for A:T>G:C, G:C>C:G, transitions, and overall intra- and interspecies variation in bias (Fig. S21). We conclude that the degree of mutational bias predicts the bias in the overall frequency of adaptive outcomes, in addition to the bias in adaptive substitution rates.

## Discussion

The hypothesis that mutational biases influence the course of adaptation predicts that species with stronger mutational biases will also exhibit stronger biases among adaptive changes. Here, we tested this prediction by compiling a dataset that, for each of 14 species, pairs previously observed adaptive amino acid substitutions with an independently measured nucleotide mutation spectrum. The strength of mutational bias for or against particular types of substitutions differed by roughly an order of magnitude across species, and these differences in mutation bias were strongly predictive of the types of mutations fixed during adaptation, producing a similar type and magnitude of bias among adaptive changes. These results suggest that interspecific differences in mutation spectrum shape adaptive outcomes in a graded manner, wherein mutational biases do not merely enrich for specific mutational classes among adaptive changes, but rather produce a bias in adaptive changes that scales positively with the bias in mutational inputs.

Existing population-genetic theory allows a range of possibilities for the extent to which mutational biases influence adaptive change. Direct proportionality is expected in the limit of low mutational supply [2], but higher levels of mutational supply can produce evolutionary dynamics that show anywhere from no relationship [12] to a strong positive relationship [17], depending on the details of the model (see Appendix A of [18]). Under some scenarios, one might even anticipate a negative relationship due to biased depletion of the distribution of fitness effects [72, 73]; that is, to the extent that early steps in an adaptive walk exhaust high-rate mutations, later steps will show a bias toward low-rate mutations. Our results suggest that adaptive evolution in the contexts examined here, specifically laboratory experiments and the natural evolution of drug resistance, mostly occurs in a regime where mutational biases do indeed have a strong influence on adaptive outcomes.

One limitation of our study is that our dataset, while unprecedented in terms of size and taxonomic breadth, is still not large enough to support analysis of more complex mutational models that include context effects, which are known to be important in work on hotspots [10, 11], cancer mutation signatures [74], and ordinary germline mutation [75, 76]. Another limitation is that our approach is based on correlations, and thus does not directly demonstrate a causal influence of mutational biases. However, the causal influence of mutational biases is already well-established, as has been shown by experimentally manipulating the mutation spectrum and observing the predicted effect on adaptation [8, 11]. In principle, an alternative explanation of our results could be that selective preferences might coincide with mutational preferences, either accidentally or due to some directional process, as argued by Shapiro [77] or Svensson and Berger [71]. Notably, however, where selection coefficients for specific adaptive changes have been measured, they do not align with mutational preferences and are not sufficient to account for the observed adaptive spectra without taking mutational biases into account ([13]; see also the analyses of [47, 15] in [3]). Finally, our source data for adaptive changes is drawn primarily from short-term microbial laboratory evolution experiments, and thus these results have no direct relevance to assessing the degree to which mutational biases shape the evolution of highly polygenic quantitative traits [78, 79].

An important area for future work is the development of additional datasets comprising measurements of both selection coefficients and mutation rates across the same set of adaptive mutations [13, 47, 15]. Knowing the joint distribution of mutation rates and selection coefficients [80] would make it possible to quantitatively partition responsibility for patterns of evolutionary change between selective preferences and mutational biases [81], providing a more granular understanding of how mutational and selective forces combine to determine the genetic basis of adaptation.

## Methods

### *De novo* mutation spectra

For 12 of the 14 species featured in our study, the spectra of *de novo* mutations were obtained from published mutation-accumulation studies. Raw mutation-accumulation data and genomic nucleotide composition are each obtained from the same study in the case of *B. cenocepacia* [20], *D. discoideum* [33], *L. lactis* [41], *P. fluorescens* [50], *P. falciparum* [48], *S. enterica* [59], and *S. aureus* [41]. For *P. aeruginosa*, raw mutation-accumulation data [46] was corrected for nucleotide composition data obtained from a separate study [82]. Raw mutation-accumulation data for *B. subtilis* [24], *C. glutamicum* [28], *E. coli* [35, 36], and *S. cerevisiae* [54, 55, 56], were each corrected for nucleotide composition obtained directly from published genome sequences: NCBI Reference Sequence NC 000964.3 (*B. subtilis*), NC 003450.3 (*C. glutamicum*), U00096.3 (*E. coli*), and GCF 000146045.2 (*S. cerevisiae*). For the remaining 2 species, *M. tuberculosis* and *T. gondii*, mutation spectra were estimated from patterns of segregating variation [12, 61], see SI Methods for details.

For each species, mutation spectra reflect raw mutation counts corrected for nucleotide composition,

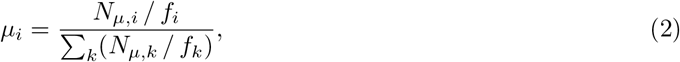

where *µ*_*i*_, *N*_*µi*_, and *f*_*i*_ are, respectively, the relative mutation rate, the raw number of observed mutations, and the relative mutational target size (the fraction of AT content in the case of mutations at A:T sites or the fraction of GC content in the case of mutations at G:C sites), for mutations of type *i*. In some cases, published mutation rates were based on counts obtained from multiple mutation-accumulation lines having different nucleotide compositions. In such cases, the calculation of *µ*_*i*_ aggregates across multiple paired values of *N*_*µ*_,*i* and *f*_*i*_ for each mutation class *i*. In order to apply pseudocounts in a consistent manner across all cases, we applied a pseudocount of *1/N*_*µ*_ to *µ*_*i*_, where *N*_*µ*_ = ∑_*k*_ *N*_*µ,k*_ is the total number of observations, and then renormalized to obtain our final *µ*_*i*_. Unless otherwise specified, *mutation bias* is defined as the relative rate of one mutation over that of others, that is, *µ*_*i*_ */* (1 − *µ*_*i*_) for mutation class *i*.

To calculate bootstrap confidence intervals for the estimated mutation biases in each species, the empirical mutation spectrum *µ*_1_, …, *µ*_6_ is treated as a multinomial distribution and is sampled *N*_*µ*_ times. The mutation spectra from which bootstrap samples were drawn do not contain pseudocounts; this was so that pseudocounts could be applied to each bootstrap sample directly. Normalizing the sample distribution results in a bootstrap mutation spectrum. We compute 95% bootstrap confidence intervals for *µ*_*i*_/(1 *− µ*_*i*_) based on 10,000 bootstrap samples.

### Spectra of adaptive changes

To reliably examine mutational influences on adaptive protein evolution, we sought to assemble a dataset that maximizes the inclusion of *bona fide* adaptive changes while minimizing the inclusion of neutral or deleterious hitchhikers. In addition, we restricted our dataset to taxa that meet two criteria. First, we only include taxa for which a sufficient number of adaptive substitution events are available to permit the estimation of an adaptive substitution spectrum. Second, we only include taxa for which an independently sourced *de novo* mutation spectrum is publicly available (see Section *De novo* mutation spectra). The resulting dataset consists of 5,488 adaptive amino acid substitutions across 14 species (Table 1). These data represent adaptive substitutions identified in *B. cenocepacia* [21, 22, 23], *B. subtilis* [25, 26, 27], *C. glutamicum* [29, 30, 31, 32], *D. discoideum* [34], *E. coli* [37, 38, 39, 40], *L. lactis* [42, 43, 44, 45], *M. tuberculosis* [12], *P. aeruginosa* [47], *P. falciparum* [49], *P. fluorescens* [51, 52], *S. aureus* [53], *S. cerevisiae* [57, 58], *S. enterica* [60], and *T. gondii* [62, 63].

To ensure that we maximized inclusion of adaptive changes while minimizing neutral mutations, we prioritized using data from experimental evolution studies, with the exception of *M. tuberculosis*, for which adaptive substitutions were obtained from published meta-analyses of antibiotic-resistance evolution [12, 64]. Amino acid substitutions identified in experimental evolution studies were evaluated on an individual basis and were included in our dataset depending on the evidence of an adaptive effect. Although the details varied by study and species, substitutions were typically only included in our study if they met both of two criteria for being adaptive. The first criterion is that substitutions were genetically linked to an adaptive phenotype, in that a newly derived allele was observed in evolved lines that express the adaptive phenotype. The second criterion is that there is some additional line of evidence implicating the substitution in the cause of the adaptive phenotype, e.g. from follow-up experiments or due to a known causal role of the affected locus, as in the case of substitutions observed to alter a known target of a drug during evolution of resistance to the same drug. See SI Methods for a detailed discussion of the adaptive substitutions dataset.

To compare biases in adaptive outcome against biases in mutation rates, raw counts of adaptive events were converted into relative rates for each class of single-nucleotide mutation. To do this, we first computed the relative frequencies of single-nucleotide nonsynonymous changes for each mutation class,

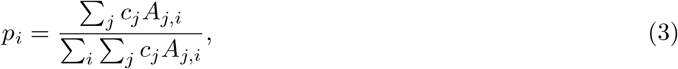

where *p*_*i*_ is the fraction of total missense paths that are accessible via a mutation of mutation class *i, c*_*j*_ is the species-specific fraction of sense codon *j* (obtained from the Codon Usage Database [83]), and *A*_*j,i*_ is the number of available missense mutations of mutation type *i* from codon *j*. While the above analysis weights all accessible nonsynonymous mutations equally, in an alternative approach that takes amino acid exchangeability into account we set *A*_*j,i*_ = Σ_*k*_ *uj*_,*k*_ where the sum is taken over all nonsynonymous codons k that are accessible from *j* by a nucleotide mutation of type *i*, and *u*_*j,k*_ is Tang’s U [69] for the pair of amino acids encoded by codons *j* and *k*.

The spectrum of adaptive changes is the vector of values computed by normalizing adaptive event counts of a given mutation class to the corresponding fraction of accessible missense paths,

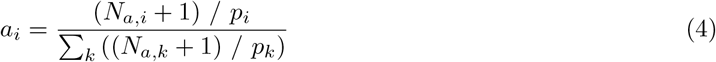

where *a*_*i*_, *N*_*a,i*_, and *p*_*i*_ are the relative rate of adaptive change, the number of observed adaptive changes, and fraction of accessible missense paths for mutations of type *i*, respectively, and the 1 is a pseudocount added to ensure that the frequency of each class of adaptive changes is positive. As with mutation bias, adaptive substitution bias is defined as the relative rate of one type of adaptive change as compared to the total of all the others, that is, *a*_i_/(1 − *a*_*i*_) for mutation class *i*. When combining multiple mutation classes, we sum their individually normalized rates rather than pooling their observed counts and mutational opportunities. This allows us to compare the same combinations of class-specific rates in the mutation and adaptive substitution spectra, without introducing different weightings due to differences in mutational opportunities among the mutation classes.

For each species, bootstrap confidence intervals on adaptive substitution bias were derived by drawing *N*_*a*_ = Σ _*k*_ *N*_*a,k*_ times with replacement from the multinomial distribution of empirical adaptive changes, that is, from the distribution of values *N*_*a,i*_/ Σ_*k*_ *N*_*a,k*_ for the six mutation classes. Each resulting dataset of bootstrap values *N*_*a,1*_,…,N_a,6_ was then used to produce a bootstrap adaptive substitution spectrum using Eq. 4. We compute the 95% bootstrap confidence intervals for *a*_*i*_/(1 − *a*_*i*_) based on 10,000 bootstrap samples.

### Statistical analysis

Interpreting Eq. 1 as a simple linear model where the log mutational bias and the log bias in adaptive outcomes are, respectively, the independent and dependent variables, we estimated the effect of mutation bias (*β*) and the intercept (*β*_0_) by ordinary least squares regression. We estimated *β* and *β*_0_ with respect to variation across taxa in transition bias, overall variation in mutation bias (within and across taxa) across all six mutation classes, and variation across taxa in mutation bias for each of the six mutation classes individually.

Uncertainty was assessed via bootstrapping and significance was assessed via comparison against simulated data under the null model in which biases in mutation rate have no effect on the adaptive outcome. To estimate bootstrap confidence intervals for the parameters of these regressions, the set of adaptive changes for each species was resampled by drawing *N*_*a*_ = Σ_*k*_ *N*_*a*,_*k* times with replacement from the multinomial distribution of empirical adaptive changes for that species. Based on each synthetic dataset, the *a*_*i*_ were then re-estimated using Eq. 4 and the regression coefficients were re-estimated. Bootstrap confidence intervals were based on 10,000 bootstrap samples.

To test the hypothesis that the magnitude of bias among adaptive outcomes depends on the strength of mutation bias, we simulated data under a null model in which biases in mutation rate have no effect on adaptive outcomes (n=10,000 simulations). Specifically, under the null model, the set of adaptive substitutions is given by sampling *N*_*a*_ times from the multinomial distribution defined by Eq. 3. The same sampling procedure was used for the analysis that corrects for amino acid exchangeability, but the sample spectra were normalized in Eq. 4 using the modified form of Eq. 3 that incorporates Tang’s U. For each simulation, we sampled a new dataset of the same size as our original data and re-estimated the parameters in the corresponding regression model. We define the p-value as the frequency of simulations that give *β*_null_ ≥ *β*_empirical_.

### Isolating the effect of interspecific differences in mutation bias

A 6-intercept model was interpreted similarly as the ordinary least squares regression model, with a single parameter *β* that captures the effect of mutation bias on bias in adaptive outcome. However, to isolate the contribution of interspecies variation in mutation bias toward *β*, while controlling for intraspecies variation, we allow the intercept *β*_*0,i*_ to be specific to mutation class *i*:

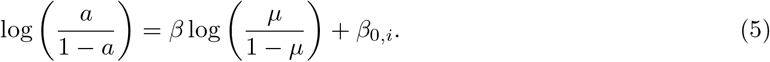

The resulting regression is fit by maximum likelihood, assuming zero-mean Gaussian errors on a log bias scale, with a single shared error variance. Statistical analysis and hypothesis-testing were performed using the same bootstraps and null-model simulations that were generated for the ordinary least squares regressions.

### Site bias and alternate-base bias

The effect of variation in site bias is assessed using the ratio of the summed rates for mutations originating at G:C sites to the corresponding sum for mutations originating at A:T sites. In particular, we perform a least-squares regression of the log ratio for adaptive substitutions against the log ratio for mutations,

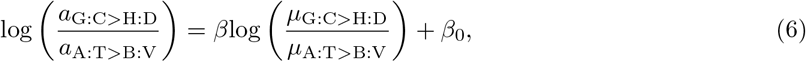

where B:V and H:D are the IUPAC representations for any base-pairs other than A:T and G:C, respectively. The quantities *a*_A:T>B:v_ and µ_A:T>B:v_ are the sums of the three components of the corresponding normalized rate spectra for mutations originating at A:T sites, while *a*_G:_c>_H:D_ and µ_G:C>H:D_ are the corresponding sums for mutations originating at G:C sites.

The effect of variation in alternate-base bias is assessed using the log bias toward each alternate base, conditioned on the identity of the wild-type base:

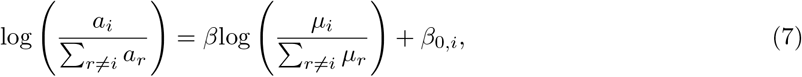

where *r* is any mutation class that shares the same wild-type base-pair with mutation class i.

### Weighted least squares regression analysis

To account for species-specific uncertainties in mutation and adaptive substitution biases, we re-estimated the effect parameter *β* for each of the regressions that were previously performed by the ordinary least squares method (see section Statistical analysis), using a weighted total least squares regression approach. To do this, we numerically minimized the expression,

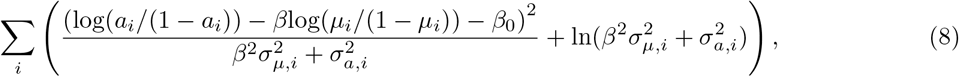

in which 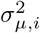 and 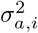 represent the variance in log mutation bias and log adaptive substitution bias, respectively, for mutation class *i*. We estimated these variances as:

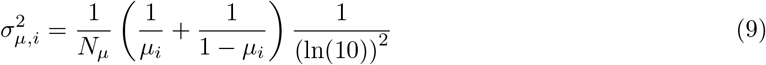

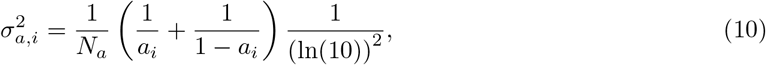

where the right-hand term accounts for the fact that we are conducting our regressions on a log base 10 scale.

To assess the impact of uncertainty in mutation bias, and variation across species in this uncertainty, we performed the weighted total least squares regression, in addition to a weighted least squares regression that only considers uncertainties in bias among adaptive outcomes. To do this, 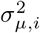 in Eq. 8 is set to zero and the resulting expression is numerically minimized with respect to the empirical data. Statistical analysis and hypothesis-testing were performed using the same bootstrap and simulated null-model datasets that were generated for the primary statistical analyses.

### Assessing the effect of neutral contamination in the adaptive substitutions data

For each species, we calculated the spectrum of adaptive substitutions under the assumption that the adaptive substitutions dataset is contaminated by a specified fraction of neutral mutations. The fraction of the adaptive dataset attributed to contamination ranged from 0 to 0.4 in increments of 0.1. Since our intention was to determine the robustness of our conclusions to the possibility of such contamination, we set the spectrum for the contamination fraction to equal the *de novo* mutation spectrum. This allows us to assess the robustness of our conclusion to a worst-case scenario in which the contamination contributes maximally toward the apparent effect of mutation bias on adaptive outcomes. We specify an expected adaptive spectrum, ϵ, given a certain fraction of contamination, *c*, which is a mixture of two component spectra, such that,

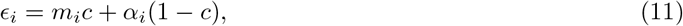

where *m*_*i*_ is the fraction of total missense mutations (see section Effect of mutation bias with respect to bias in frequencies of adaptive outcomes) and α_*i*_ is the fraction of adaptive substitutions not attributed to contamination, respectively, for mutation class *i*. We numerically calculated α as the frequency distribution that minimizes the sum of squared differences between ϵ and the empirical adaptive frequency spectrum, *N*_*a,i*_/*N*_*a*_. We multiplied α by *N*_*a*_ (1 − c) to obtain inferred uncontaminated adaptive event counts. For rate-based analyses, these counts were subsequently converted to adaptive substitution rates using Eq. 4. We then repeated the regression analyses and simulations described in the previous sections, using the biases among inferred contamination-free adaptive substitutions.

### Effect of mutation bias with respect to bias in frequencies of adaptive outcomes

To assess the effect of mutation bias on the relative frequencies with which adaptive outcomes are observed, we consider the spectrum of raw adaptive changes, that is, without normalizing the adaptive counts of a given mutation class to the corresponding fraction of accessible missense paths:

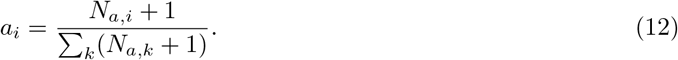

The frequency spectrum among *de novo* missense mutations was estimated as

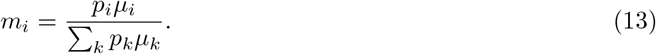

We then repeated the ordinary least squares and 6-intercept model regression analyses as before, using biases in the frequency with which mutation classes are represented among *de novo* missense mutations versus adaptive outcomes, with the same bootstrapping and null-model simulation procedures that were described previously.

## Data, Materials, and Software Availability

Model fitting, bootstrapping, and maximum-likelihood inference were conducted using a custom Python script. Code and source data are available at https://github.com/bgitschlag/mbamta.

## Acknowledgments

This work was made possible through the support of the John Templeton Foundation (grant #61782, D.M.M.). The opinions expressed in this publication are those of the authors and do not necessarily reflect the views of the John Templeton Foundation. D.M.M. also acknowledges additional support from the Simons Center for Quantitative Biology.

## Data availability

The data and code used in this work are available from a GitHub repository ( HYPERLINK “https://github.com/bgitschlag/mbamta”https://github.com/bgitschlag/mbamta HYPERLINK “https://github.com/bgitschlag/mbamta”).

## Notes

https://github.com/bgitschlag/mbamta

### Competing Interest Statement

The authors have declared no competing interest.

### Summary of Updates

The manuscript and analysis pipeline have been updated to add robustness tests and to revise the main analysis to define and compute biases in mutation rates and adaptive substitution rates, rather than frequencies as in the initial submission. These revisions were performed in response to reviewer comments as part of the preparation of the manuscript for resubmission to a journal.

