## Supporting Information for "Molecular adaptation reflects taxon-specific mutational biases"

**This PDF file includes:**

Supporting Methods
Figures S1 to S21
Tables S1 to S8
SI References

### SI Methods: inference of mutation spectra

In addition to mutation-accumulation studies, mutation spectra for 2 of the 14 species are derived from polymorphism data. In particular, the mutation spectrum for *M. tuberculosis* was obtained in a previous study, in which synonymous single-nucleotide polymorphisms (SNPs) in third codon positions were corrected for total nucleotide composition across third codon positions [1]. Finally, to derive the raw mutation spectrum for *T. gondii*, we utilized published SNP data [2]. Based on the reasoning that SNPs are less likely to be affected by selection if they are recently derived, in noncoding regions, or both, we restricted our estimation of the mutation spectrum to only include SNPs in noncoding regions that occurred recently enough to be unique to one of the three sub-strains derived from the RH laboratory strain (see Figure 3B in [2]; "non-coding" and "intron" mutations that are unique to B-RH, G-RH, or 2F-1 were included). To correct for nucleotide composition, we computed the nucleotide composition of the noncoding regions in *T. gondii*. To do this, the genome-wide nucleotide composition (0.520 GC) and the fraction of the genome size that represents coding sequences (0.279) were obtained from a single study [3]. The GC content of the coding regions (0.5795, obtained from the Codon Usage Database [4]) was weighted by the relative size of the coding regions. Subtracting the size-weighted GC content from the genome-wide content and normalizing to the size of the noncoding fraction of the genome gives the nucleotide composition (0.497 GC) of the noncoding fraction.

### SI Methods: assembly of the adaptive substitutions dataset

**Bacillus subtilis.** Adaptive changes in *B. subtilis* were obtained from three published experimental evolution studies. The first study used experimental evolution to identify mutations in *B. subtilis* that enhance colonization of plant roots [5], using an experimental design that involves growth of *B. subtilis* populations on *Arabidopsis thaliana* root, tomato root, or alternation between Arabidopsis and tomato roots. Genome sequencing revealed 14 nonsynonymous mutations that appeared and rose to high frequency during adaptation to plant roots, to the point that they had become the majority alleles by the end of the experiment. We include mutations in our dataset of adaptive changes if they meet at least two separate criteria for having an adaptive effect. Specifically, in addition to having risen to high allele frequency, mutations were included in our dataset if they altered a protein with a function previously associated with plant root colonization, namely cell motility or biofilm production [6, 7, 8]. Notably, these account for the majority (9 of the 14) of the mutations that rose to high frequency during adaptation. Secondly, we also included mutations as adaptive if, in addition to having risen to high frequency, they did so in at least two independent lineages. Taken together, 12 nonsynonymous mutations meet these criteria for adaptive effects.

A second study likewise used experimental evolution to recover mutations associated with enhanced colonization of tomato plant roots, alone as well as in the presence of a competing bacterial species, namely *Pseudomonas fluorescens* [9]. Although this study did not report allele frequencies for the mutations, several nonsynonymous mutations were observed to be present in consecutive timepoints (that is, the mutation does not appear and then disappear), including the endpoint of the evolution experiment. We included such mutations in our dataset of adaptive changes if they occurred in a gene known to have a phenotypically relevant effect within the context of the evolution experiment. For example, the gene pksR was altered during adaptation to the presence of *P. fluorescens* but not in the absence of *P. fluorescens*. [9] attribute adaptation in the pksR-mutated lineage to a deletion in the gene ywcC, since the latter mutation contributes most strongly toward root colonization. However, we note a modest adaptive benefit at a timepoint in which only the pksR mutation is reportedly present (see timepoint C7 in Figure 3B of [9]). Although this effect was not reported to be statistically significant, we include the pksR mutation as adaptive due to a qualitative judgment that this timepoint shows higher root colonization, as well as the fact that the gene pksR is involved in the synthesis of the antibiotic bacillaene, previously identified to inhibit the growth of other bacteria in the genus *Pseudomonas* [10]. In total, 5 nonsynonymous mutations meet these criteria for adaptive effects.

The third study used experimental evolution to recover mutations associated with adaptation of *B. subtilis* to the presence of the fungus *Aspergillus niger* [11]. As in the case of the study by Hu, *et al.* [5], genome sequencing of consecutive time points revealed several nonsynonymous changes that arose spontaneously and reached high frequency during adaptive evolution, becoming the majority alleles by the end of the experiment. Interestingly, four of the five control lineages, propagated in the absence of *A. niger*, were reported to phenotypically resemble biofilm matrix over-producers, a previously reported adaptive phenotype [8, 12]. Accordingly, our adaptive dataset includes mutations from both the coevolved and control conditions, provided the mutations meet at least two criteria for having an adaptive effect. Applying a similar standard of evidence as for the two previous studies, we identified mutations as adaptive if, in addition to having risen to high frequency, the mutations were phenotypically characterized in follow-up experiments to show an adaptive effect. The mutations in the genes degU and degS meet these criteria, in particular. Finally, mutations were also included as adaptive if, in addition to having achieved high frequency during adaptive evolution, they occurred in pathways or macromolecular complexes that were observably mutated in independently evolving lineages. In addition to the 17 mutations from the previous studies, 12 additional adaptive changes meet these criteria for an adaptive effect, resulting in a combined dataset of 29 individual adaptive changes in *B. subtilis*.

**Burkholderia cenocepacia.** Adaptive changes in *B. cenocepacia* were obtained from three published experimental evolution studies. One study grew *B. cenocepacia* under conditions that select for biofilm-mediated growth on a hard surface, by selectively propagating bacteria that had become attached to polystyrene beads in liquid medium [13]. Beginning with these evolved biofilm specialists, the second study sought to identify mutations that restore fitness under selection for planktonic growth in liquid suspension, in the absence of polystyrene beads [14]. Numerous nonsynonymous point mutations were reported to arise during adaptation to these specific niches. We included such mutations in our dataset of adaptive changes if, in addition to these adaptive circumstances, they met at least two criteria for having an adaptive effect. Specifically, we only considered mutations as adaptive if they were associated with an adaptive phenotype under the specific selection conditions. These consist of the "wrinkly" colony morphology in the case of biofilm-mediated growth on hard surfaces, or a wrinkly-suppressed "smooth" colony morphology in the case of selection for planktonic growth. Consistent with a second criterion of adaptive effect, such mutations were reported to occur either in genes with a known biofilm-mediated colony phenotype, namely the "wrinkly spreader" wsp genes, or in a locus that was found to be mutated in multiple parallel evolving lineages [14], in a manner that cannot be attributed to a mutation hotspot since the specific mutations differ between lineages. Of the changes that met these criteria, 30 missense changes can be unambiguously attributed to specific single-nucleotide mutations, given the structure of the genetic code, under the assumption that multi-nucleotide mutations are rare.

The third study sought to explore the evolvability of antibiotic resistance in the presence of an adjuvant [15], under conditions that select for resistance either in the presence of antibiotic alone or the antibiotic together with the adjuvant baicalin hydrate (BH). This study identified 20 independent events consisting of unambiguous single-nucleotide mutations, 12 of which occurred in antibiotic-exposed populations but were absent in antibiotic-free control populations. Of the 12 single-nucleotide mutations that occurred exclusively in antibiotic-exposed populations, 11 were nonsynonymous, resulting in either the replacement of an amino acid with another (6 mutations) or the introduction of a stop codon (5 mutations). In addition to having arisen exclusively during adaptation to the presence of antibiotic, with or without BH, all 11 mutations meet at least one further criterion for having an adaptive effect. For example, the C131W change in CepI arose exclusively in the presence of antibiotic in 3 lineages, 2 of which occurred in the presence of BH. This change was shown to confer BH resistance biochemically, by increasing the amount of BH needed to inhibit CepI enzymatic activity. Moreover, we include the amino acid change F51L, which occurred in an ABC-type transporter protein, in our study since this mutation occurred in a carbohydrate transporter and rose to fixation during evolution of resistance to a glycoside antibiotic, namely tobramycin. The remaining 7 mutations were found to affect either a gene or a biochemical process that was altered in independently evolving populations. These include 5 mutations in an ABC peptide transporter, which was found to be altered in 3 separate ways across 5 of the 6 antibiotic-exposed populations, in addition to 2 mutations in malate dehydrogenase. Interestingly, malate dehydrogenase catalyzes the formation of oxaloacetate, a function that is shared by the enzyme phosphoenolpyruvate carboxylase, which was also mutated during evolution of resistance to tobramycin and BH. Taken together, we identified 6 adaptive amino acid substitutions during evolution of tobramycin resistance, with or without BH, resulting in a combined dataset of 36 adaptive missense changes when we include those that occurred during adaptation to surface-attached versus planktonic growth.

**Corynebacterium glutamicum.** Adaptive changes in *C. glutamicum* were obtained from four published studies, including three experimental evolution studies and one study that focused on characterizing the consequences of deleting the metabolic enzyme aconitase [16, 17, 18, 19]. Deletion of aconitase compromises the ability to convert citrate to isocitrate, which was hypothesized to select for mutations that reduce the activity of citrate synthase, the enzyme that catalyzes the previous step in the tricarboxylic acid (TCA) cycle [19]. Consistent with this hypothesis, mutations in the citrate synthase gene gltA were identified in 16 of the 28 aconitase deletion clones examined. Importantly, citrate synthase activity was reported to be lower in all 16 gltA mutants, but not in the other clones. Five of the 16 gltA mutations were single-nucleotide missense changes, which are included as adaptive in our study due to their phenotypic effect and the phenotypically relevant circumstances in which they arose. Deletion of isocitrate dehydrogenase, which catalyzes the TCA cycle step that immediately follows the aconitase-catalyzed step, was found to have a similar effect, resulting in the identification of a sixth adaptive substitution event in gltA.

One of the three experimental evolution studies focused on identifying mutations that confer higher growth rates in glucose minimal medium [16]. Multiple nonsynonymous point mutations were observed in the genes fruK and pyk during adaptation to glucose minimal medium. These genes encode phosphofructokinase and pyruvate kinase, respectively. Five unique mutational paths, representing 9 mutational events in the evolution experiment, were observed in fruK and pyk and were individually characterized via reintroduction to the wild-type strain. Although these mutations did not always rise to high frequency, remaining the minor allele in the population for 4 of the mutational events, we reason that this likely reflects clonal interference due to the presence of other adaptive changes in the populations, for two reasons. First, in the same populations where these mutations occur at low frequency, other mutations are present in at least one of these same two genes, in some cases at high frequency (see Figure 4 in [16]). Second, the individually characterized mutational paths were each shown in follow-up experiments to confer an increase in growth rate.

The two remaining studies used experimental evolution to identify mutations that enhance thermal and methanol tolerance in *C. glutamicum* [17, 18]. Two mutations were individually found to confer thermotolerance [17], and 6 were found to increase growth in up to 4 g/L methanol (see Figure 4a in [18]), 5 of which were missense changes. Together with the previous two studies, we identified 22 individual adaptive changes in *C. glutamicum*.

**Dictyostelium discoideum.** Adaptive changes in *D. discoideum* were obtained from the one known study in which *D. discoideum* genomes were sequenced and nucleotide changes were reported following experimental evolution [20]. Under nutritionally permissive conditions, *D. discoideum* are free-living amoeboid protists. Upon encountering starvation conditions, however, *D. discoideum* aggregate into a multicellular phase, in which reproduction consists of a division of labor whereby some cells form spores at the top of a fruiting body while other cells altruistically form the stalk of the fruiting body [21, 22]. Importantly, these cooperative interactions are understood to require high relatedness among cells, since low relatedness can leave the multicellular group vulnerable to cheating behaviors, in which some genotypes maximize their own inclusion among the spores, at the expense of alternative genotypes that are relegated to the non-reproductive stalks [22, 23]. Consistent with expectation, evolution under conditions that permit low relatedness resulted in the frequent loss, among parallel lineages, of cooperative fruiting body formation [24]. These evolved lines were sequenced in a more recent study, which revealed 31 single-nucleotide mutations within coding regions, distributed across 17 evolved lines [20]. Although only 21 of these mutations occurred within the coding regions of non-fruiting clones in particular, we reason that most of the 31 mutations are adaptive, for three reasons. First, adaptive evolution under conditions of low relatedness can proceed without necessarily resulting in the loss of fruiting body formation, as illustrated via antagonistic coevolution between *D. discoideum* strains [25]. Second, and more importantly, the coding-region mutations consisted of an astonishing 30:1 ratio of nonsynonymous to synonymous mutations, strongly indicative of positive selection. Finally, the 30 nonsynonymous mutations were estimated to all have moderate to high impact, according to Ensembl Variant Effect Predictor, a genomic variant annotation tool that predicts mutation impacts using the GENCODE and RefSeq gene sets [26]. Consistent with this, we note that 4 of the 8 *de novo* stop codons had occurred in the same gene that represented 6 of the 22 missense mutations, suggesting that protein alterations at this locus were under selection. Given the neutral expectation, the observed ratio of nonsynonymous to synonymous mutations suggests that approximately 90 percent (27 of 30) of the nonsynonymous changes are *bona fide* drivers of adaptation. Accordingly, we include the 22 missense changes in our dataset of adaptive substitutions. To account for the possibility of non-adaptive contaminants in these data, we later assess the robustness of our overall conclusions to the contamination of our dataset due to the inclusion of non-adaptive mutations (see section Assessing the effect of non-adaptive contamination).

**Escherichia coli.** Adaptive changes in *E. coli* were obtained from four published experimental evolution studies, which were identified for their evidence of individual mutations having adaptive effects. Two studies selecting for improved growth on minimal media identified 15 nonsynonymous single-nucleotide mutations, 4 of which confer statistically significant growth-rate increases on glycerol minimal medium [27] with the remaining 11 mutations conferring significantly elevated growth rates on lactate minimal medium (see Table 2 in [28]). Two additional studies selected for growth at elevated temperature [29, 30]. From these thermal adaptation studies, we included mutations as adaptive changes if they met at least one of two criteria of evidence for an adaptive effect. First, adaptation to high temperature involved a substantial number of alterations to proteins which have been independently associated with tolerance to physiological stress, such as temperature and osmotic stress. These include gene expression machinery, namely RNA polymerase subunits [31, 32, 30] and transcriptional regulators such as sigma and termination factors [31, 33], in addition to the cardiolipin synthetase Cls [34, 35], the ion antiporter YbaL [31], and cell wall proteins MrdA and MrdB [36]. Notably, although mutations of the isoleucine at position 15 in the transcriptional termination factor Rho are deleterious on their own [37], we include them as adaptive for three reasons. Specifically, such mutations were reported by Tenaillon *et al.* [29] to have occurred no less than 17 times in independently evolving lines, suggesting that these changes likely confer an adaptive benefit under certain conditions, such as in combination with other mutations. Moreover, consistent with this possibility, mutation of I15 in Rho was always accompanied by alterations to at least one, and usually two, of the proteins YbaL, Cls, or the transcriptional regulator IclR. Lastly, further consistent with epistatic effects, mutations of I15 in Rho were more recently shown to be associated with a fitness gain when in combination with mutations in the RNA polymerase gene rpoB [38]. In total, 282 single-nucleotide missense changes reported in [29] met these criteria.

In addition to the mutations identified by [29] to occur in known stress-tolerance genes during adaptation to high temperature, we included mutations in our dataset of adaptive changes if they were obtained during adaptation to high temperature and were individually shown to confer a statistically significant fitness increase at 42.4°C [30]. Four additional missense mutations met these criteria, 2 of which occurred in RNA polymerase subunits, consistent with the findings reported by [29]. Taken together, a total of 301 mutations in *E. coli* occurred during adaptive evolution and met our criteria for evidence of an adaptive effect.

**Lactococcus lactis.** Adaptive changes in *L. lactis* were obtained from four published experimental evolution studies. These studies focused on identifying mutations that contribute toward adaptation to aerated growth conditions [39] and high temperature [40, 41], as well as mutations that confer adaptation of a plant-associated strain to a dairy niche [42]. One of the three dairy-adapted strains reportedly harbored more than twice as many mutations as the other two strains combined, including a deletion in a DNA repair gene [42], and was therefore identified as a mutator strain. We omitted any changes in the mutator strain from our adaptive dataset, due to the possibility that its mutation spectrum has diverged from that of other *L. lactis* strains. From the remaining strains across all four studies, we included any missense mutations as adaptive changes in our study if they arose spontaneously during the adaptation experiments and met at least one further criterion for having an adaptive effect. One such criterion is parallel change. In particular, mutations are more likely to be adaptive if, in addition to having been observed to arise during adaptation, they constitute multiple nonsynonymous changes in the same gene or in the same general biological process, such as nucleotide metabolism. Additionally, we include a given mutation as an adaptive change if it affects a gene or process that is phenotypically relevant under the experimental conditions; that is, a gene or process that has been associated in the literature with a phenotype that affects survival or growth under the conditions in which the mutation arose. A final criterion of adaptive change consists of functional characterization, in which the mutations were empirically shown to contribute to the adaptive phenotype. We identify 28 missense mutations meeting such criteria for evidence of an adaptive effect in *L. lactis*.

**Mycobacterium tuberculosis.** Adaptive changes in *M. tuberculosis* were obtained from a prior study, which explored the influence of mutation spectra on adaptive outcomes within species [1]. The study merged the resulting datasets from two previously published meta-analyses [43, 44], which focused on mutations in *M. tuberculosis* that drive adaptation to antibiotic stress. Mutational paths consisted of those that passed stringent criteria of evidence for resistance-conferring effects, and the number of independent mutational events for each mutational path was inferred from phylogenetic analysis using publicly available M. tuberculosis genome sequencing data [1]. Merging the two datasets resulted in 4,414 adaptive events across 256 missense mutational paths (see Dataset S3 in ref. [1]).

**Plasmodium falciparum.** Adaptive changes in *P. falciparum* were obtained from a study that sought to identify antimalarial drug targets and drug-resistance mechanisms, using an experimental evolution approach that selected for resistance to a library of compounds [45]. The study identified numerous target genes that met one of two criteria for being involved in resistance evolution. First, consistent with expectation, some genes that were mutated during adaptation to a particular compound were previously associated with drug resistance. Alternatively, some genes were mutated during adaptation to numerous compounds independently. Following similar reasoning, we included any additional genes from the original study if the encoded protein was altered during the independent evolution of resistance to different compounds. Finally, applying a further criterion for evidence of adaptive effects, we include missense mutations in our dataset as adaptive changes if they occurred in one of these genes and if they were not accompanied by synonymous mutations in the same gene. This criterion was based on the reasoning that, since adaptive protein evolution tends to increase the ratio of nonsynonymous to synonymous changes, the observation of exclusively nonsynonymous mutations at a given locus raises the likelihood that the nonsynonymous changes have an adaptive effect. A total of 84 missense changes passed our criteria for adaptive effects.

**Pseudomonas aeruginosa.** Adaptive changes in *P. aeruginosa* were obtained from a study that explored the evolution of resistance to the antibiotic rifampicin [46]. After a period of selecting for rifampicin resistance by cultivating *P. aeruginosa* in the presence of rifampicin, sequencing was performed to identify mutations in the gene rpoB, which encodes the RNA polymerase $\beta$ subunit and target of rifampicin [47]. Importantly, since the study focused on epistatic effects that occur when resistance mutations arise in different ancestral backgrounds, an empirical fitness assay was used to compare the fitness of evolved versus ancestral strains. Accordingly, all mutations from the *P. aeruginosa* study included in our adaptive dataset were confirmed to confer a fitness advantage over their respective ancestral genotypes. A total of 144 adaptive missense events were observed.

**Pseudomonas fluorescens.** Adaptive changes in *P. fluorescens* were obtained from two previously published experimental evolution studies that explored the genetic basis of adaptive divergence into niche specialists [48, 49]. In spatially structured microcosms consisting of static liquid nutrient broth in glass vials, *P. fluorescens* readily evolve to occupy the air-liquid interface, either stably or transiently, corresponding to phenotypes labeled wrinkly spreader (WS) or fuzzy spreader (FS), respectively, named after their colony morphologies [50, 49]. Mutations included as adaptive changes in our dataset were single-nucleotide missense changes that arose during adaptation to the structured microcosms and were individually shown to confer the WS phenotype [48] or the FS phenotype [49]. Notably, the V148G amino acid change occurred 28 times, accounting for more than half of all single-nucleotide missense changes that were observed in the evolution of FS or FS-like phenotypes [49], suggesting that the underlying nucleotide change (T443G) represents a mutation hotspot. To minimize the confounding influence of hotspot mutations in our analysis, the V148G events were omitted from our adaptation dataset. After omitting these hotspot changes, a remaining total of 35 adaptive missense changes in *P. fluorescens* were identified and included in our study.

**Saccharomyces cerevisiae.** Adaptive changes in *S. cerevisiae* were obtained from two published experimental evolution studies, which were chosen for their large numbers of mutations that meet strong evidence of adaptive effects. The first study selected for resistance of *S. cerevisiae* populations to individual xenobiotic compounds, in parallel evolution experiments across a library of 80 compounds [51]. Sequencing revealed 1,405 mutations across 355 compound-resistant clones. From these data, we sought to maximize the inclusion of adaptive driver mutations in our study whilst minimizing the inclusion of non-adaptive mutations. To this end, we filtered these data to include only single-nucleotide mutations that meet at least one of three specific criteria for evidence of an adaptive effect. First, a subset of the nonsynonymous mutations reported in [51] appeared in the activation domains of proteins that were previously associated with multidrug resistance and which were repeatedly altered in the evolution of resistance to a wide range of compounds. In particular, the genes yrm1 and yrr1, encoding transcriptional regulators of multidrug resistance [52, 53, 54], were mutated on at least 83 independent occasions spanning the evolution of resistance to 19 compounds, in which all such mutations were nonsynonymous and occurred in the C-terminal activation domains of YRM1 and YRR1. In addition, several mutations either altered the target protein of a compound during the evolution of resistance to the same compound, or occurred in a protein that was previously associated with resistance to the same compound. For example, the most commonly altered protein during the evolution of resistance to the topoisomerase poison etoposide was the Type II DNA topoisomerase TOP2, whereas the most commonly altered protein during evolution of resistance to the drug cipargamin was PMA1, which had previously been implicated in cipargamin resistance [55]. Finally, a subset of the mutations were individually shown to confer resistance to the compounds that were used in the evolution condition in which the same mutations initially arose. Of the 1,405 mutations in the original dataset, a total of 130 single-nucleotide missense mutations met at least one of these three criteria and were thus included in our study as adaptive changes.

The second study reported 43 yeast clones that had acquired heightened tolerance to caffeine during experimental evolution [56]. Following a similar standard of evidence as in [51], we included mutations in our dataset of adaptive changes if they arose during evolution of caffeine tolerance and they nonsynonymously altered a target gene, defined as a gene that was either previously associated with caffeine tolerance or was shown by [56] to affect caffeine sensitivity. Strikingly, all 43 clones carried at least 1 mutation, but fewer than 2 on average, in a target gene. Moreover, the target-gene mutations were single-nucleotide nonsynonymous changes in 42 of the 43 clones. In total, 72 such missense mutations were identified. Taken together with the mutations reported in [51], we identified 202 mutations in *S. cerevisiae* that meet our criteria of evidence for adaptive effects.

**Salmonella enterica (serovar typhimurium).** Adaptive changes in *S. enterica* were obtained from a study that used experimental evolution to identify compensatory mutations in streptomycin-resistant bacteria [57]. Streptomycin works by targeting the bacterial ribosome. The amino acid substitution K42N in the ribosomal protein S12 confers streptomycin resistance at the expense of a reduced bacterial growth rate. Adaptive changes included in our study consist of missense mutations in ribosomal proteins that were identified as mutations that restore bacterial fitness [57]. These evolution experiments were performed in four ancestral genetic backgrounds, three of which are suspected to have different mutation spectra due to the presence of mutations that affect genes involved in either DNA replication or DNA repair. We therefore only include adaptive changes that arose during evolution from an ancestral strain that carried the K42N mutation in an otherwise wild-type genetic background. A total of 30 adaptive amino acid changes were attributable to specific nucleotide changes and were therefore included in our study as adaptive events.

**Staphylococcus aureus.** Adaptive changes in *S. aureus* were obtained from a study that used experimental evolution to identify mutations that confer resistance to the antibiotic ciprofloxacin [58], which targets bacterial DNA topoisomerase enzymes. After culturing *S. aureus* populations in the presence of ciprofloxacin, sequencing of newly-resistant populations uncovered 104 single-nucleotide missense changes in one of the DNA topoisomerase genes, which were therefore included in our dataset as adaptive changes.

**Toxoplasma gondii.** Adaptive changes in *T. gondii* were obtained from two experimental evolution studies. One study focused on causal mutations for resistance to the antimalarial artemisinin [59], while the other sought to identify compensatory mutations that reverse the fitness cost of mutations that confer resistance to the herbicide oryzalin [60]. In the case of both studies, we include mutations in our dataset of adaptive changes if they were phenotypically characterized and individually shown to confer a survival or growth rate advantage in comparison to their respective genetic background. This corresponds to 4 unique missense mutations in two genes, in the case of artemisinin resistance [59]. With respect to the evolutionary recovery of fitness following acquisition of oryzalin resistance, we include mutations as adaptive changes in our study if they were shown to at least partially restore fitness in their respective oryzalin-resistant genetic backgrounds, without entirely eliminating oryzalin resistance (see Table 3 in [60]; mutations are included as adaptive if the decrease in resistance and the population decrease are $< 100\%$). A total of 37 amino acid changes pass these criteria and can be assigned an unambiguous single-nucleotide mutation based on the reference codon and the amino acid change.

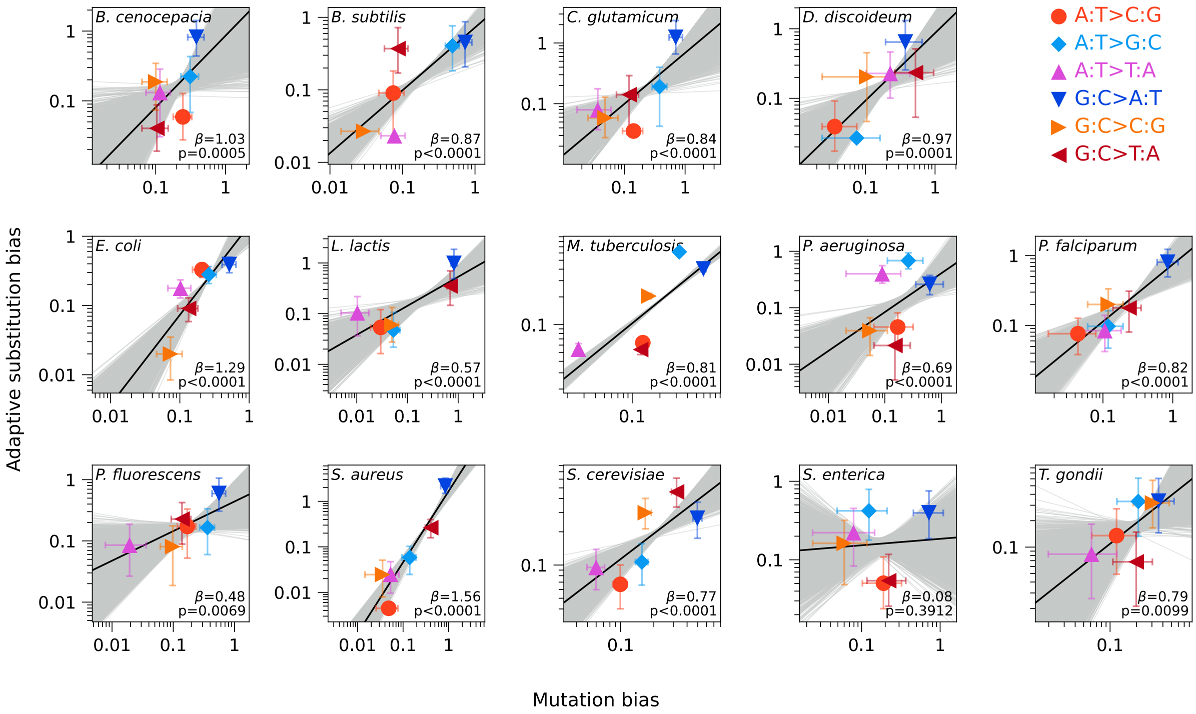


Fig. S1: **Biases in the spectrum of adaptive substitutions reflect biases in the mutation spectrum, in 13 of 14 species.** Bias among adaptive substitutions, for each of the six mutation classes, is plotted as a function of mutation bias. Biases are measured as rate ratios for each of the mutation classes. Least squares regression on log-transformed data yields a significantly positive slope for 13 of 14 species. Regressions on empirical data are shown with solid black line and the 10,000 bootstrap regressions are shown with gray lines. Significance was assessed by simulating datasets (n=10,000) under the null model in which biases in mutation rate have no effect on the adaptive outcome. See Table S1 for summary statistics.

| Species | $\beta$ | $\beta_{\text{null}}$ | Pearson's $r$ | p |
|---|---|---|---|---|
| *B. cenocepacia* | 1.03 (0.41–1.70) | -0.67–0.65 | 0.58 (0.23–0.79) | 0.0005 |
| *B. subtilis* | 0.87 (0.63–1.08) | -0.36–0.37 | 0.78 (0.57–0.89) | <0.0001 |
| *C. glutamicum* | 0.84 (0.48–1.22) | -0.42–0.43 | 0.77 (0.47–0.89) | <0.0001 |
| *D. discoideum* | 0.97 (0.55–1.35) | -0.40–0.51 | 0.82 (0.48–0.94) | 0.0001 |
| *E. coli* | 1.29 (1.04–1.66) | -0.22–0.22 | 0.82 (0.73–0.88) | <0.0001 |
| *L. lactis* | 0.57 (0.37–0.83) | -0.25–0.27 | 0.83 (0.58–0.94) | <0.0001 |
| *M. tuberculosis* | 0.81 (0.76–0.87) | -0.04–0.04 | 0.79 (0.76–0.82) | <0.0001 |
| *P. aeruginosa* | 0.69 (0.43–1.01) | -0.25–0.25 | 0.41 (0.25–0.55) | <0.0001 |
| *P. falciparum* | 0.82 (0.56–1.09) | -0.31–0.31 | 0.92 (0.71–0.98) | <0.0001 |
| *P. fluorescens* | 0.48 (0.15–0.91) | -0.35–0.37 | 0.78 (0.25–0.95) | 0.0069 |
| *S. aureus* | 1.56 (1.39–1.82) | -0.22–0.22 | 0.93 (0.87–0.98) | <0.0001 |
| *S. cerevisiae* | 0.77 (0.53–1.02) | -0.24–0.25 | 0.77 (0.56–0.88) | <0.0001 |
| *S. enterica* | 0.08 (-0.37–0.53) | -0.52–0.49 | 0.08 (-0.32–0.45) | 0.3912 |
| *T. gondii* | 0.79 (0.22–1.53) | -0.60–0.64 | 0.70 (0.19–0.92) | 0.0099 |

Table S1: **Summary statistics for intraspecific regressions.** Summary statistics are shown for least squares regression of log biases in the adaptive substitution spectrum against log biases in the mutation spectrum, for each of the 14 species (Fig. S1). We report the statistic $\beta$ that gives the slope of the regression line, capturing the effect of mutation bias on biases in the rates of adaptive substitutions, as well as Pearson's correlation coefficient. For $\beta_{\text{null}}$, the range corresponds to 95% of the slopes calculated when simulating data under the null model where mutation bias has no effect on bias in adaptive substitutions (n=10,000 simulated datasets). Columns for $\beta$ and Pearson's $r$ show estimates for the empirical data with 95% bootstrap confidence intervals in parentheses, and the p-value is given by the fraction of simulated datasets under the null where the slope equals or exceeds the slope calculated from the empirical data.

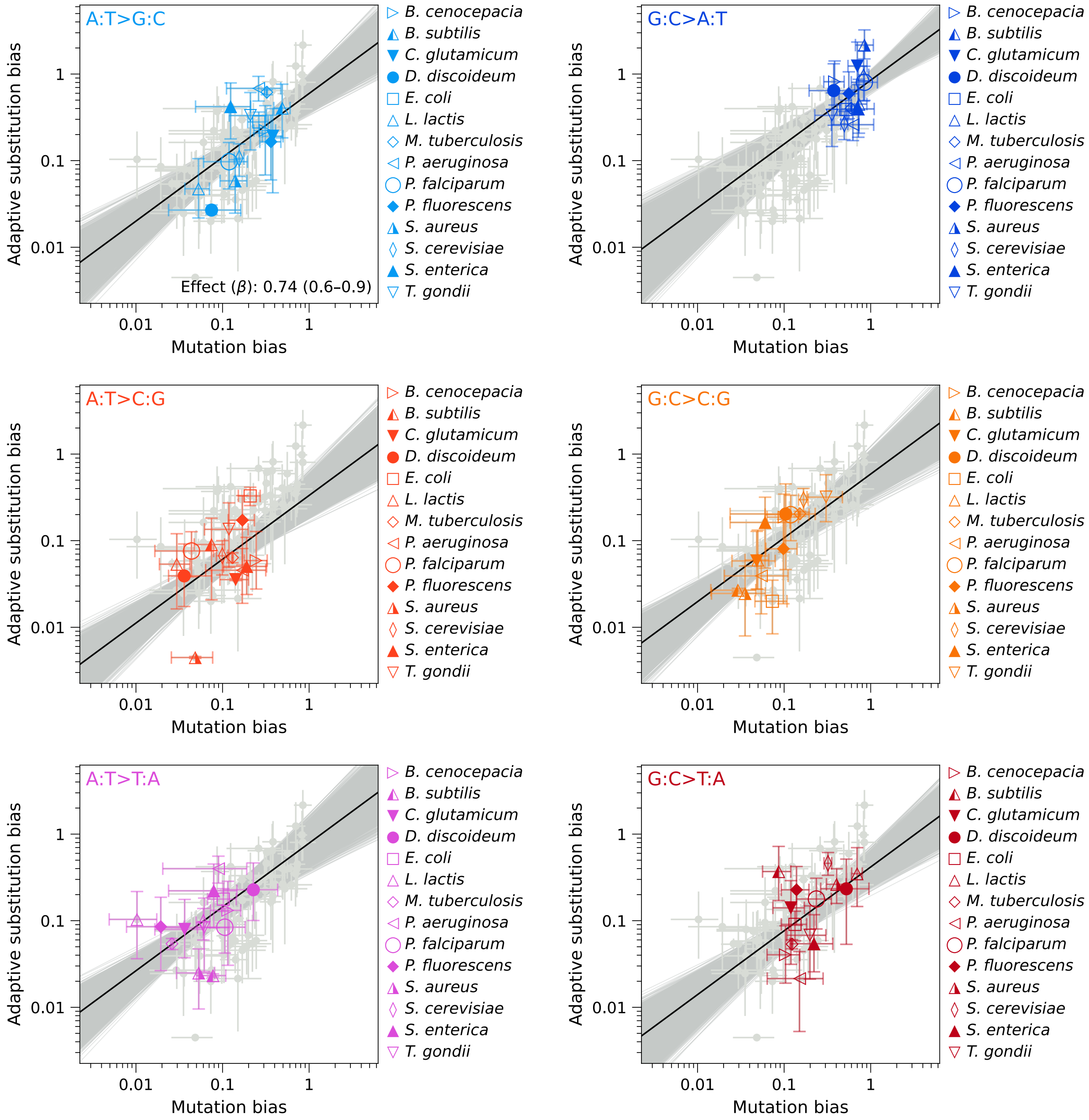


Fig. S2: **Variation across species in overall bias in adaptive substitution spectra reflects variation in bias in mutation spectra.** Bias among adaptive substitutions is plotted against mutation bias, across 14 species, for each of the six single-nucleotide mutation classes. The regression analysis shown here isolates interspecies variation using a single mutation bias effect parameter $\beta$ influenced by all mutation classes, along with class-specific intercepts, on log-transformed empirical data (solid black line) and 10,000 bootstrap datasets (gray lines). The effect parameter $\beta$ is shown in the plot for A:T>G:C (upper left) with 95% bootstrap confidence interval in parentheses. Significance was assessed by simulating datasets (n=10,000) under the null model in which mutation bias has no effect on bias in adaptive substitutions, $p<0.0001$.

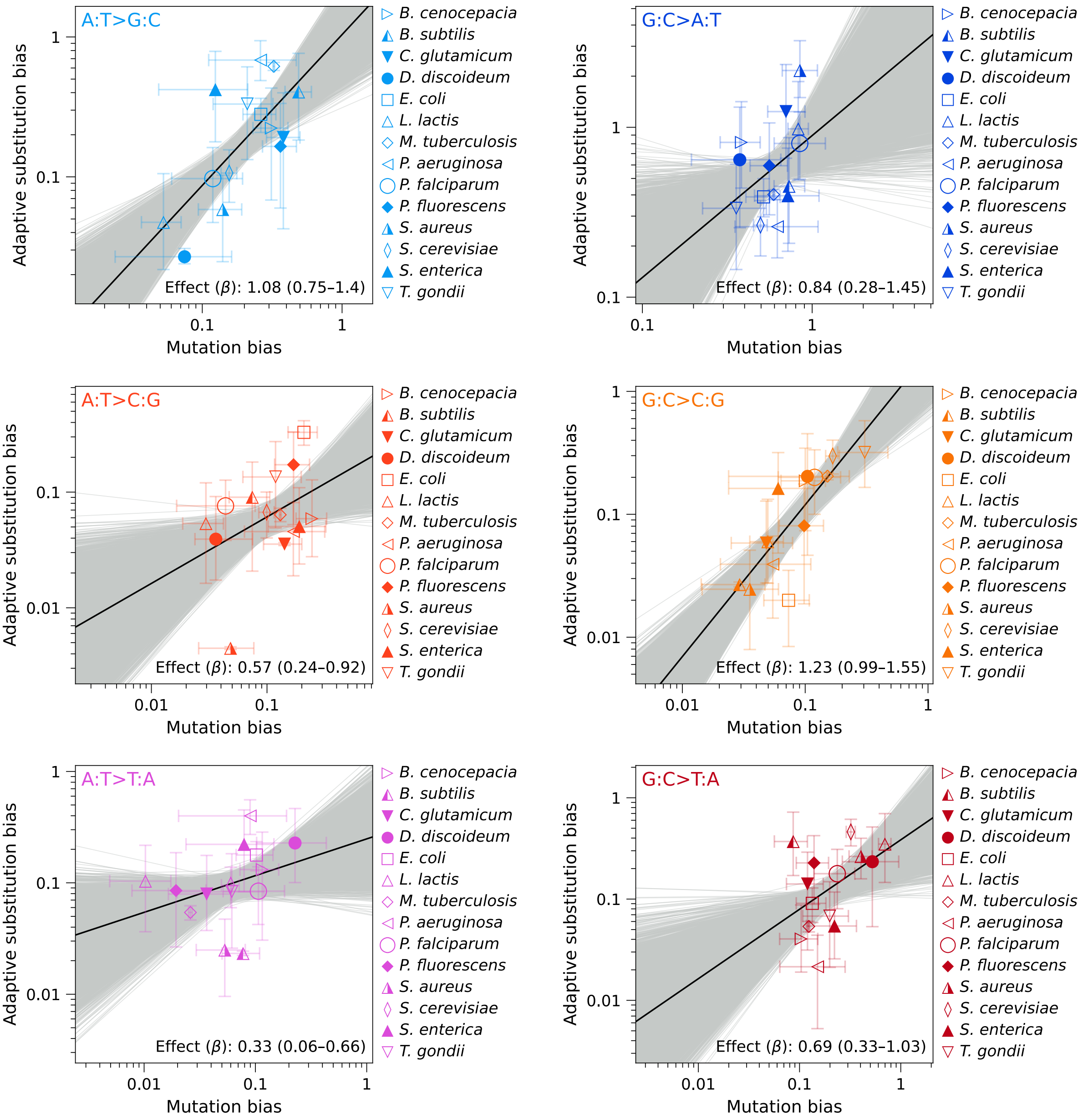


Fig. S3: **Variation across species in adaptive substitution bias reflects variation in mutation bias, for each of the single-nucleotide mutation classes.** Biases in the rates of adaptive substitutions are plotted against mutation bias, across 14 species. The data and regression analyses (solid black lines) are the same as in Fig. 5, but separated by mutation class into individual plots, with bootstrap regressions (n=10,000) shown as gray lines. The 95% bootstrap confidence interval for mutation bias effect, $\beta$, is shown in parentheses for each regression. Significance (shown in Fig. 5 and Table S2) was assessed by simulating datasets (n=10,000) under the null model in which mutation bias has no effect on bias in adaptive substitutions, $p \leq 0.022$ for all slopes.

| | Combined (intra- & interspecies) and mutation class-specific effects | | | | | Isolation of interspecies variation in bias, aggregating across the six mutation classes | | | |
|---|---|---|---|---|---|---|---|---|---|
| Mutation or bias type | $\beta$ | $\beta_{\text{null}}$ | $\beta_0$ | Pearson's $r$ | p | $\beta$ | $\beta_{\text{null}}$ | $\beta_0$ | p |
| Transition | 0.67 (0.28–1.11) | -0.43–0.43 | 0.04 (-0.03–0.12) | 0.60 (0.24–0.76) | 0.0012 | N/A | N/A | N/A | N/A |
| Intra- & inter-species | 0.80 (0.73–0.91) | -0.08–0.11 | -0.20 (-0.27– -0.14) | 0.70 (0.61–0.73) | <0.0001 | N/A | N/A | N/A | N/A |
| A:T>C:G | 0.57 (0.24–0.92) | -0.25–0.34 | -0.64 (-1.00– -0.32) | 0.41 (0.16–0.60) | 0.0006 | 0.74 (0.60–0.90) | -0.13–0.17 | -0.48 (-0.65– -0.32) | <0.0001 |
| A:T>G:C | 1.08 (0.75–1.40) | -0.33–0.40 | 0.02 (-0.26–0.24) | 0.72 (0.48–0.82) | <0.0001 | | | -0.22 (-0.36– -0.10) | |
| A:T>T:A | 0.33 (0.06–0.66) | -0.27–0.27 | -0.61 (-0.95– -0.23) | 0.33 (0.06–0.56) | 0.0109 | | | -0.10 (-0.30–0.08) | |
| G:C>A:T | 0.84 (0.28–1.45) | -0.76–0.81 | -0.05 (-0.20–0.09) | 0.41 (0.12–0.61) | 0.0220 | | | -0.07 (-0.15– -0.0020) | |
| G:C>C:G | 1.23 (0.99–1.55) | -0.30–0.32 | 0.31 (0.02–0.61) | 0.82 (0.64–0.88) | <0.0001 | | | -0.23 (-0.41– -0.06) | |
| G:C>T:A | 0.69 (0.33–1.03) | -0.37–0.39 | -0.42 (-0.71– -0.19) | 0.46 (0.21–0.61) | 0.0002 | | | -0.38 (-0.53– -0.26) | |

Table S2: **Summary statistics on key results.** Summary statistics are shown for least squares regression analyses from Figs. 3 to 5 and Fig. S2. Regression analyses were performed on log-transformed data. We report the statistic $\beta$ that gives the slope of the regression line, capturing the effect of mutation bias on biases in the rates of adaptive substitutions, as well as Pearson's correlation coefficient. For $\beta_{\text{null}}$, the range corresponds to 95% of the slopes calculated when simulating data under the null model where mutation bias has no effect on bias in adaptive substitutions (n=10,000 simulated datasets). Columns for $\beta$ and Pearson's $r$ show estimates for the empirical data with 95% bootstrap confidence intervals in parentheses, and the p-value is given by the fraction of simulated datasets under the null where the slope equals or exceeds the slope calculated from the empirical data. The lefthand set of columns gives the results for Figs. 3 to 5 and the righthand set of columns gives the results for Fig. S2 (where the 6 mutational classes each have their own intercept $\beta_{0,i}$ but share a single slope $\beta$).

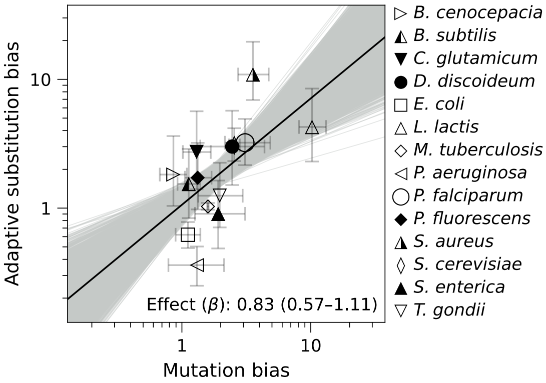


Fig. S4: **Mutational site bias contributes to site bias in adaptive substitutions.** Site bias in rates of adaptive substitutions is plotted against site bias in *de novo* mutation rates, using least squares regression on the log-transformed empirical data (solid black line) and 10,000 bootstrap datasets (gray lines). The 95% bootstrap confidence interval for mutation bias effect, $\beta$, is shown in parentheses. Significance is assessed by simulating datasets (n=10,000) under the null model in which mutation bias has no effect on bias in adaptive substitutions, $p<0.0001$.

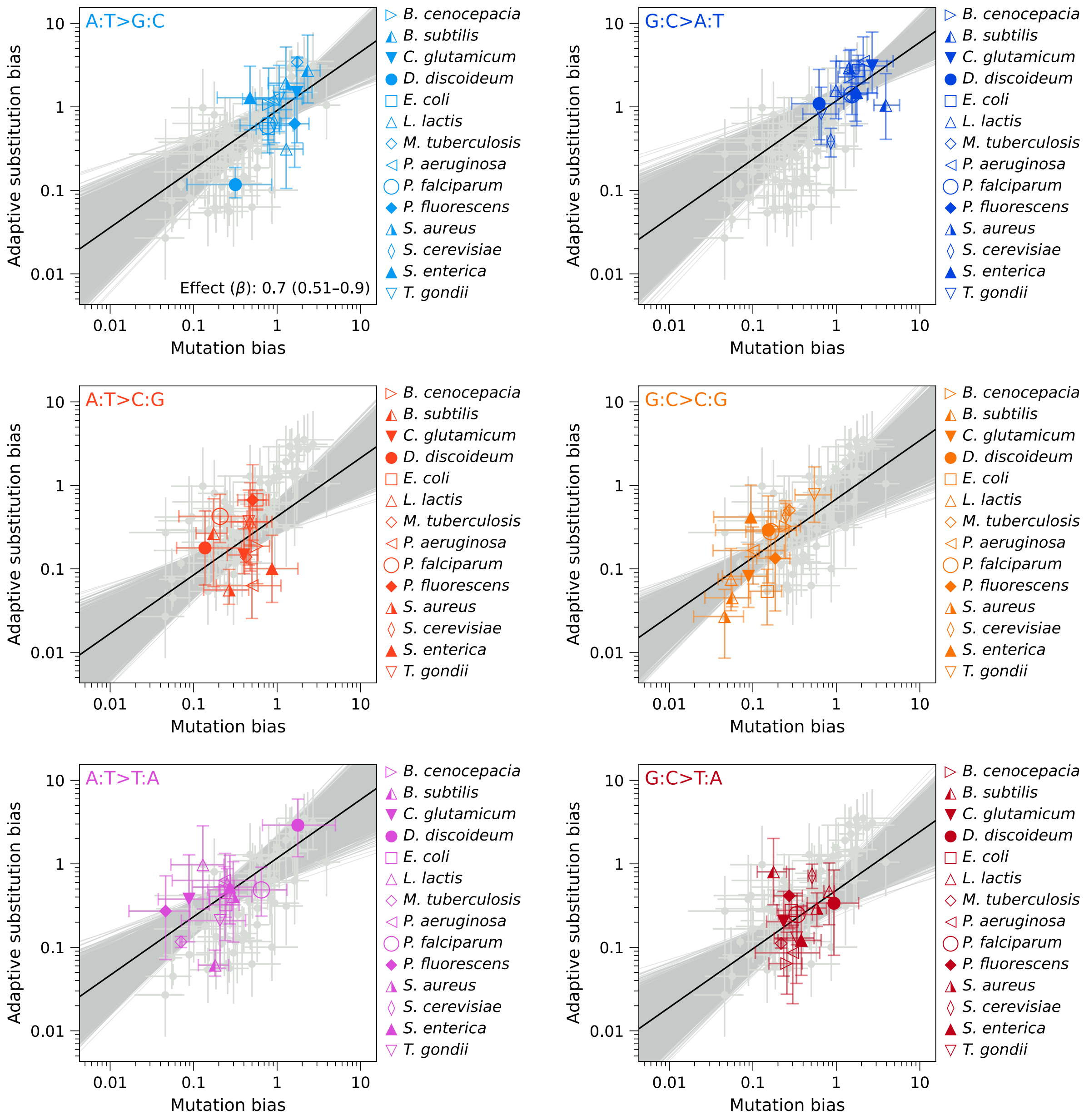


Fig. S5: **Variation in alternate-base bias contributes strongly toward the effect of mutation bias on adaptive substitutions.** Bias among adaptive substitutions within a species is plotted as a function of the corresponding mutation bias, across 14 species, for each of the six single-nucleotide mutation classes. This is similar to Fig. S2 but plotted values represent bias between the alternate (mutant allele) bases, conditioned on the identity of the mutated reference base, so that the magnitude of bias is unaffected by variation in underlying base mutability (Fig. S4, see Methods for details). Regression analysis was performed using a single mutation bias effect parameter, $\beta$, and class-specific intercepts, on log-transformed empirical data (solid black line) and 10,000 bootstrap datasets (gray lines). The effect parameter $\beta$ is shown in the plot for A:T>G:C (upper left) with 95% bootstrap confidence interval in parentheses. Significance was assessed by simulating datasets (n=10,000) under the null model in which mutation bias has no effect on bias in adaptive substitutions, p<0.0001.

| Held-out species | Transition | Intra- & interspecies | Interspecies | A:T>C:G | A:T>G:C | A:T>T:A | G:C>A:T | G:C>C:G | G:C>T:A |
|---|---|---|---|---|---|---|---|---|---|
| *B. cenocepacia* | 0.697 | 0.795 | 0.756 | 0.672 | 1.105 | 0.323 | 1.167 | 1.217 | 0.581 |
| *B. subtilis* | 0.919 | 0.792 | 0.793 | 0.603 | 1.114 | 0.384 | 0.923 | 1.241 | 1.019 |
| *C. glutamicum* | 0.588 | 0.796 | 0.762 | 0.620 | 1.169 | 0.327 | 0.734 | 1.255 | 0.736 |
| *D. discoideum* | 0.670 | 0.787 | 0.718 | 0.605 | 0.883 | 0.239 | 1.083 | 1.211 | 0.699 |
| *E. coli* | 0.665 | 0.784 | 0.695 | 0.402 | 1.071 | 0.299 | 0.803 | 1.205 | 0.680 |
| *L. lactis* | 0.673 | 0.876 | 0.854 | 0.732 | 1.109 | 0.560 | 0.749 | 1.253 | 0.629 |
| *M. tuberculosis* | 0.637 | 0.798 | 0.707 | 0.580 | 1.007 | 0.294 | 0.839 | 1.232 | 0.629 |
| *P. aeruginosa* | 0.661 | 0.807 | 0.700 | 0.630 | 1.020 | 0.260 | 0.880 | 1.205 | 0.589 |
| *P. falciparum* | 0.670 | 0.798 | 0.760 | 0.685 | 1.070 | 0.357 | 0.823 | 1.212 | 0.678 |
| *P. fluorescens* | 0.717 | 0.830 | 0.777 | 0.506 | 1.174 | 0.369 | 0.839 | 1.248 | 0.750 |
| *S. aureus* | 0.636 | 0.703 | 0.628 | 0.277 | 1.028 | 0.313 | 0.452 | 1.178 | 0.647 |
| *S. cerevisiae* | 0.530 | 0.799 | 0.700 | 0.575 | 1.066 | 0.330 | 0.736 | 1.191 | 0.590 |
| *S. enterica* | 0.675 | 0.835 | 0.779 | 0.640 | 1.202 | 0.302 | 0.939 | 1.291 | 0.703 |
| *T. gondii* | 0.683 | 0.798 | 0.716 | 0.555 | 1.071 | 0.331 | 0.762 | 1.393 | 0.684 |

Table S3: **Robustness of results to exclusion of taxa.** Values of $\beta$ plotted in Fig. 6B are given here, along with (in column 4) values of $\beta$ from the same iterative leave-one-out approach applied to the regression that isolates total interspecies variation across all single-nucleotide mutation classes (Fig. S2), to assess the impact of omitting individual species on the estimated effect of mutation bias, $\beta$, for each of the regression analyses.

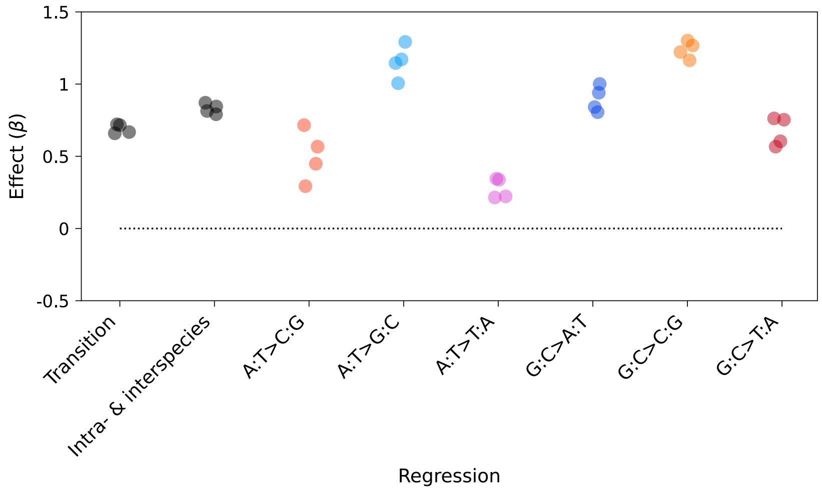


Fig. S6: **Effect of excluding specific pairs of taxa.** Shown here are values of $\beta$ upon repeating the regression analyses reported in (from left to right) Fig. 3, Fig. 4, and the individual regressions from Fig. 5 and Fig. S3, while breaking up the two closest pairs of species (Table S4), to enhance phylogenetic independence.

| Held-out species | Transition | Intra- & interspecies | Interspecies | A:T>C:G | A:T>G:C | A:T>T:A | G:C>A:T | G:C>C:G | G:C>T:A |
|---|---|---|---|---|---|---|---|---|---|
| *E. coli, P. aeruginosa* | 0.659 | 0.791 | 0.647 | 0.448 | 1.006 | 0.215 | 0.841 | 1.165 | 0.567 |
| *E. coli, P. fluorescens* | 0.716 | 0.814 | 0.731 | 0.293 | 1.171 | 0.338 | 0.806 | 1.222 | 0.753 |
| *S. enterica, P. aeruginosa* | 0.668 | 0.845 | 0.744 | 0.716 | 1.146 | 0.222 | 1.001 | 1.267 | 0.604 |
| *S. enterica, P. fluorescens* | 0.722 | 0.871 | 0.820 | 0.567 | 1.292 | 0.345 | 0.940 | 1.301 | 0.762 |

Table S4: **Robustness of results to exclusion of specific pairs of taxa.** Values of $\beta$ plotted in Fig. S6 are shown here, along with (in column 4) values of $\beta$ from the regression that isolates total interspecies variation across all single-nucleotide mutation classes (Fig. S2), to assess the impact of omitting taxa to enhance phylogenetic independence.

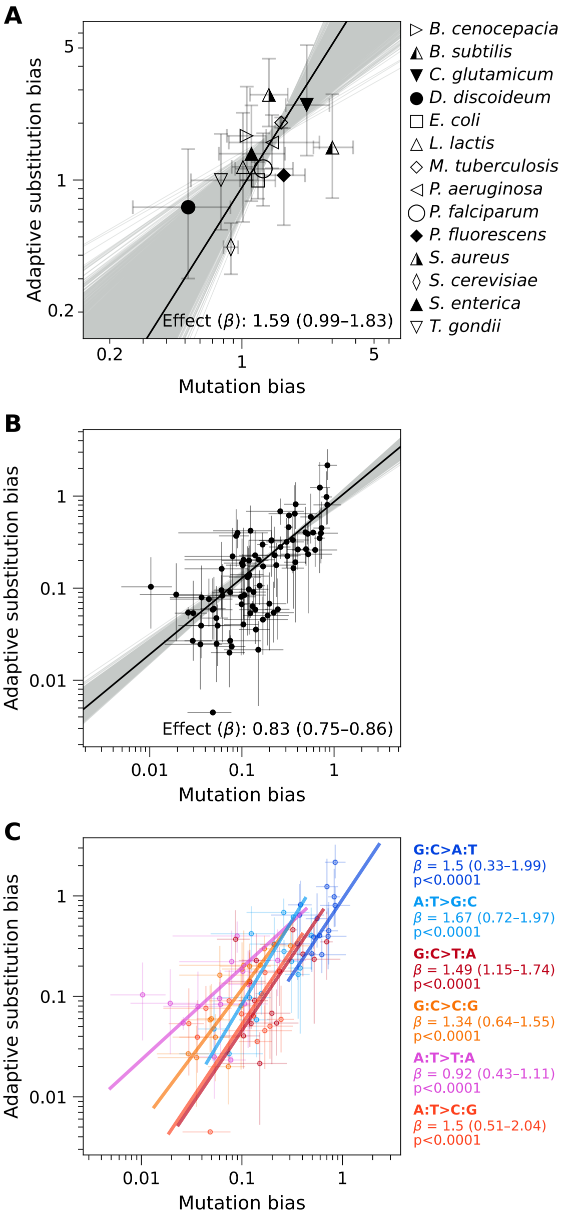


Fig. S7: **Robustness of results under weighted least squares.** Regression analyses for transition bias (Fig. 3), overall bias across taxa and mutational classes (Fig. 4), and each mutation class individually (Fig. 5) are repeated in panels A, B, and C, respectively, using a weighted least squares regression approach that uses inverse variance in adaptive substitution bias as weights. Regressions are on log-transformed empirical data (solid black line for panels A and B, and color-coded lines by mutation class for panel C), and on 10,000 bootstrap datasets (gray lines in panels A and B; not shown in panel C). The effect of mutation bias, $\beta$, is shown for each regression, with 95% bootstrap confidence interval in parentheses. Significance was assessed by simulating datasets (n=10,000) under the null model in which mutation bias has no effect on bias in adaptive substitutions ($p<0.0001$ for panels A and B, and for individual mutation class-specific p-values shown in panel C).

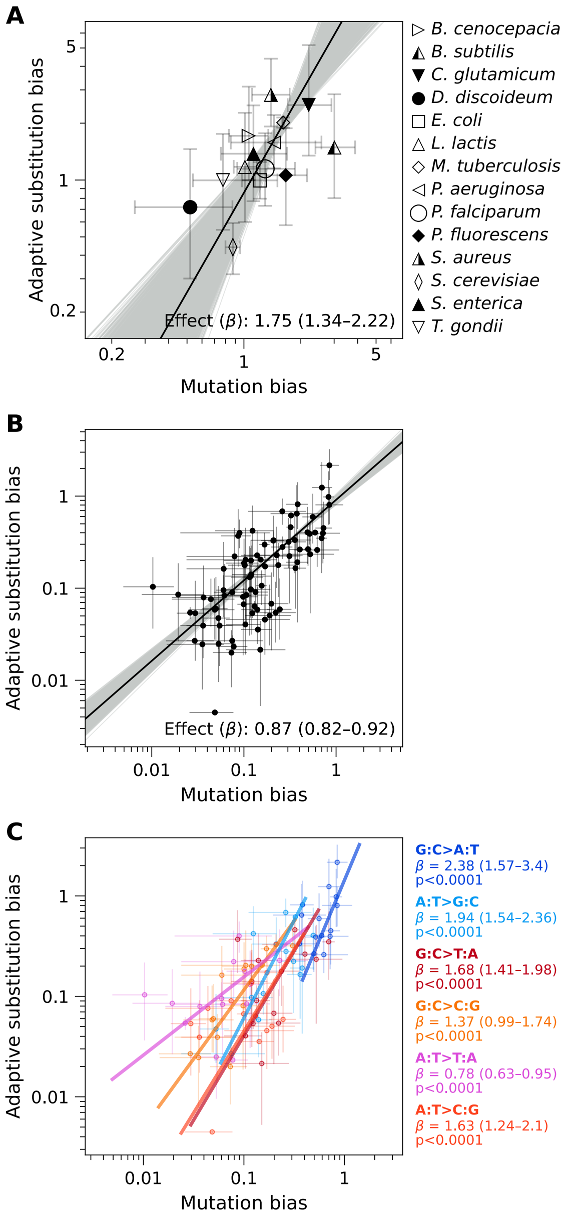


Fig. S8: **Robustness of results under weighted total least squares.** Regression analyses for transition bias (Fig. 3), overall bias across taxa and mutational classes (Fig. 4), and each mutation class individually (Fig. 5) are repeated in panels A, B, and C, respectively, similar to Fig. S7 but using a *total* weighted least squares regression approach that uses inverse variance in adaptive substitution bias and mutation bias as weights. Regressions are on log-transformed empirical data (solid black line for panels A,B and color-coded lines by mutation class for panel C), and on 10,000 bootstrap datasets (gray lines in panels A,B; not shown in panel C). The effect of mutation bias, $\beta$, is shown for each regression, with the 95% bootstrap confidence interval in parentheses. Significance was assessed by simulating datasets (n=10,000) under the null model in which mutation bias has no effect on bias in adaptive substitutions ($p<0.0001$ for panels A and B, and for individual mutation class-specific p-values shown in panel C).

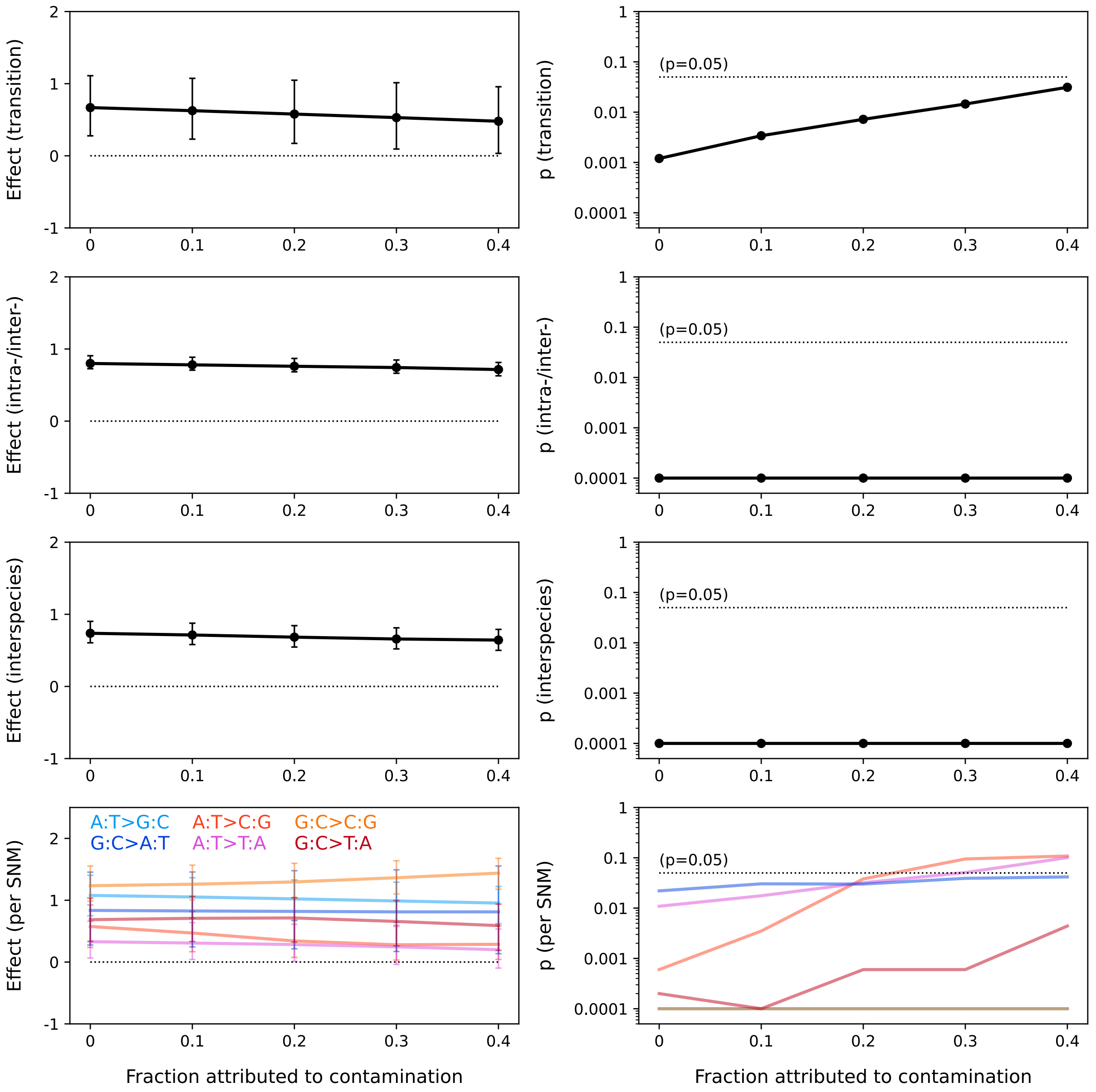


Fig. S9: **Robustness of results to contamination of the adaptive substitutions dataset.** Shown here are values of the statistic characterizing the effect of mutation bias, $\beta$, on adaptive outcome (left column) and the corresponding p-values (right column), upon increasing attribution of the adaptive substitutions dataset to non-adaptive contamination, under the assumption that the contamination spectrum (that is, the distribution of non-adaptive contaminants across the mutation classes) matches the *de novo* mutation spectrum. Regressions included in this analysis (by row, from top): transition bias (Fig. 3), combined effect of intra- and inter-species variation in bias for all single-nucleotide mutation classes (Fig. 4), the combined effect of interspecies variation across all six mutation classes (Fig. S2), and the regressions on interspecies variation for each of the six mutation classes individually (Fig. 5 and Fig. S3). Error bars show 95% bootstrap confidence intervals. Significance was assessed by simulating datasets (n=10,000) under the null model in which mutation bias has no effect on bias in adaptive substitutions.

| Mutation or bias type | $\beta$ | $p$ |
|---|---|---|
| Transition | 0.67 (0.28–1.11) | 0.0001 |
| Intra- & inter-species | 0.8 (0.73–0.91) | <0.0001 |
| Interspecies | 0.74 (0.6–0.9) | <0.0001 |
| A:T>C:G | 0.57 (0.24–0.92) | 0.0001 |
| A:T>G:C | 1.08 (0.75–1.4) | <0.0001 |
| A:T>T:A | 0.33 (0.06–0.66) | 0.0073 |
| G:C>A:T | 0.84 (0.28–1.45) | 0.0017 |
| G:C>C:G | 1.23 (0.99–1.55) | <0.0001 |
| G:C>T:A | 0.69 (0.33–1.03) | 0.0002 |

Table S5: **Alternative hypothesis test against $\beta = 0$.** Results reflect the same regression of adaptive substitution biases on mutation biases as shown in Figs. 3 to 5, Fig. S2 and Table S2, but we replaced our specific null hypothesis that the adaptive substitutions are multinomial draws from Eq. 3 with the broader and more expressive null hypothesis that $\beta = 0$. We conduct a one-sided test of this broader null by inverting our percentile bootstrap confidence intervals for $\beta$ so that the p-value is given by the fraction of the n=10,000 bootstrap samples where $\beta \leq 0$. The table reports these p-values together with the point estimates and 95% bootstrap confidence intervals from Table S2.

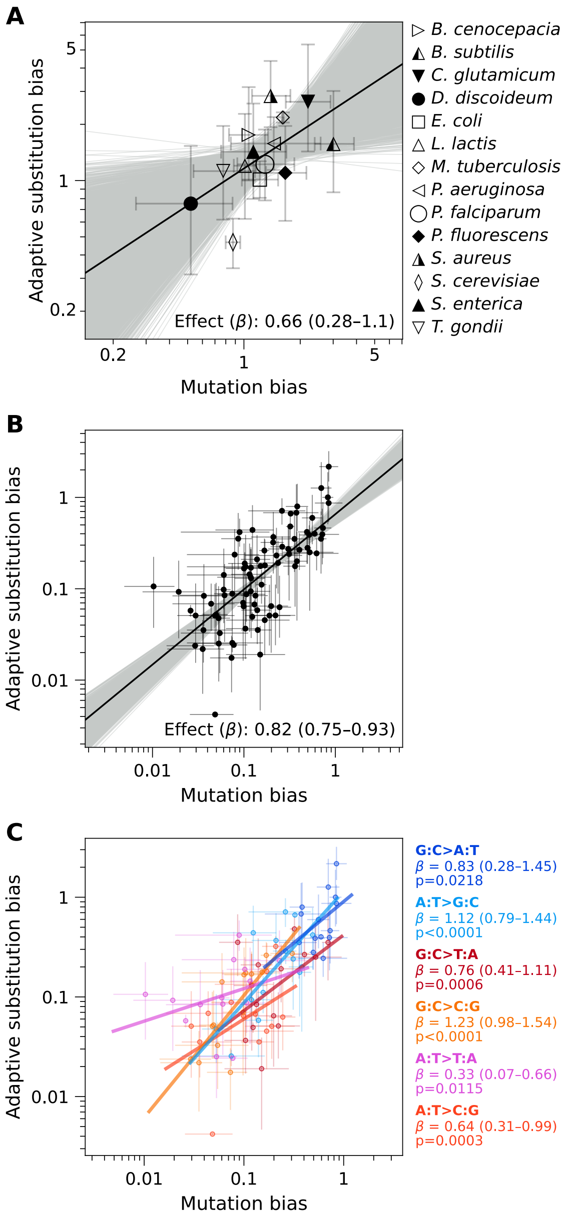


Fig. S10: **Correcting for selection due to amino acid exchangeability has a negligible effect on the association between mutation bias and bias in adaptive substitutions.** Regression analyses for transition bias (Fig. 3), overall bias across taxa and mutational classes (Fig. 4), and each mutation class individually (Fig. 5) are repeated in panels A, B, and C, respectively, using biases in adaptive substitutions that were corrected for amino acid exchangeability indices (see Methods). Least squares regressions are on log-transformed empirical data (solid black line for panels A and B, and color-coded lines by mutation class for panel C), and on 10,000 bootstrap datasets (gray lines in panels A and B; not shown in panel C). The effect of mutation bias, $\beta$, is shown for each regression, with the 95% bootstrap confidence interval in parentheses. Significance was assessed by simulating datasets (n=10,000) under the null model in which mutation bias has no effect on bias in adaptive substitutions (p=0.0014 for panel A, p<0.0001 for panel B; individual mutation class-specific p-values are shown in panel C).

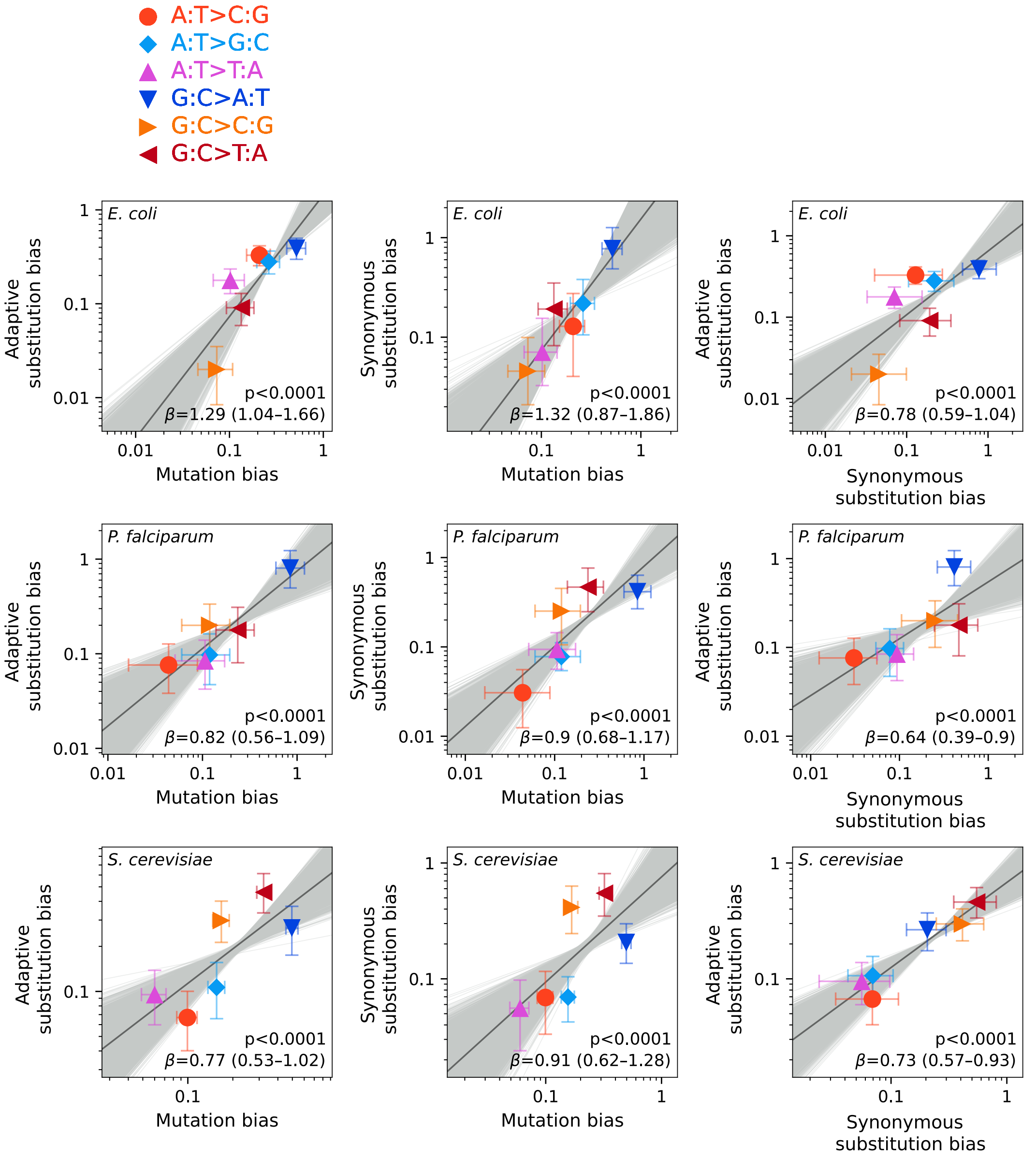


Fig. S11: **Biases in synonymous substitutions, adaptive missense substitutions, and *de novo* mutations are all positively associated with one another.** Synonymous substitutions were obtained from the same studies that served as the source literature for adaptive substitutions (see Table 1 and Methods) and normalized to the fraction of accessible synonymous mutations based on species-specific codon usage, to obtain rates of synonymous substitutions in a similar manner to the derivation of adaptive substitution rates from adaptive substitution counts. The first column shows the positive association between adaptive missense substitution bias and *de novo* mutation bias (reproduced here from the regressions shown in Fig. S1 for *E. coli*, *P. falciparum*, and *S. cerevisiae*). The middle column shows the positive association between synonymous substitution bias and mutation bias, in agreement with the association between adaptive substitution bias and mutation bias. The right column shows the association of adaptive (missense) substitution bias with synonymous substitution bias. The solid black lines show least squares regressions on log-transformed data, with gray lines showing regressions on 10,000 bootstrap datasets of adaptive substitution spectra (left and right columns) or synonymous substitution spectra (middle column). The effect of mutation bias, $\beta$, is shown for each plot in the left and middle columns. For the right column, synonymous substitution bias is treated as a proxy for mutation bias. The 95% bootstrap confidence intervals for $\beta$ are shown in parentheses. Significance was assessed by simulating datasets (n=10,000) under a null model in which mutation bias has no effect.

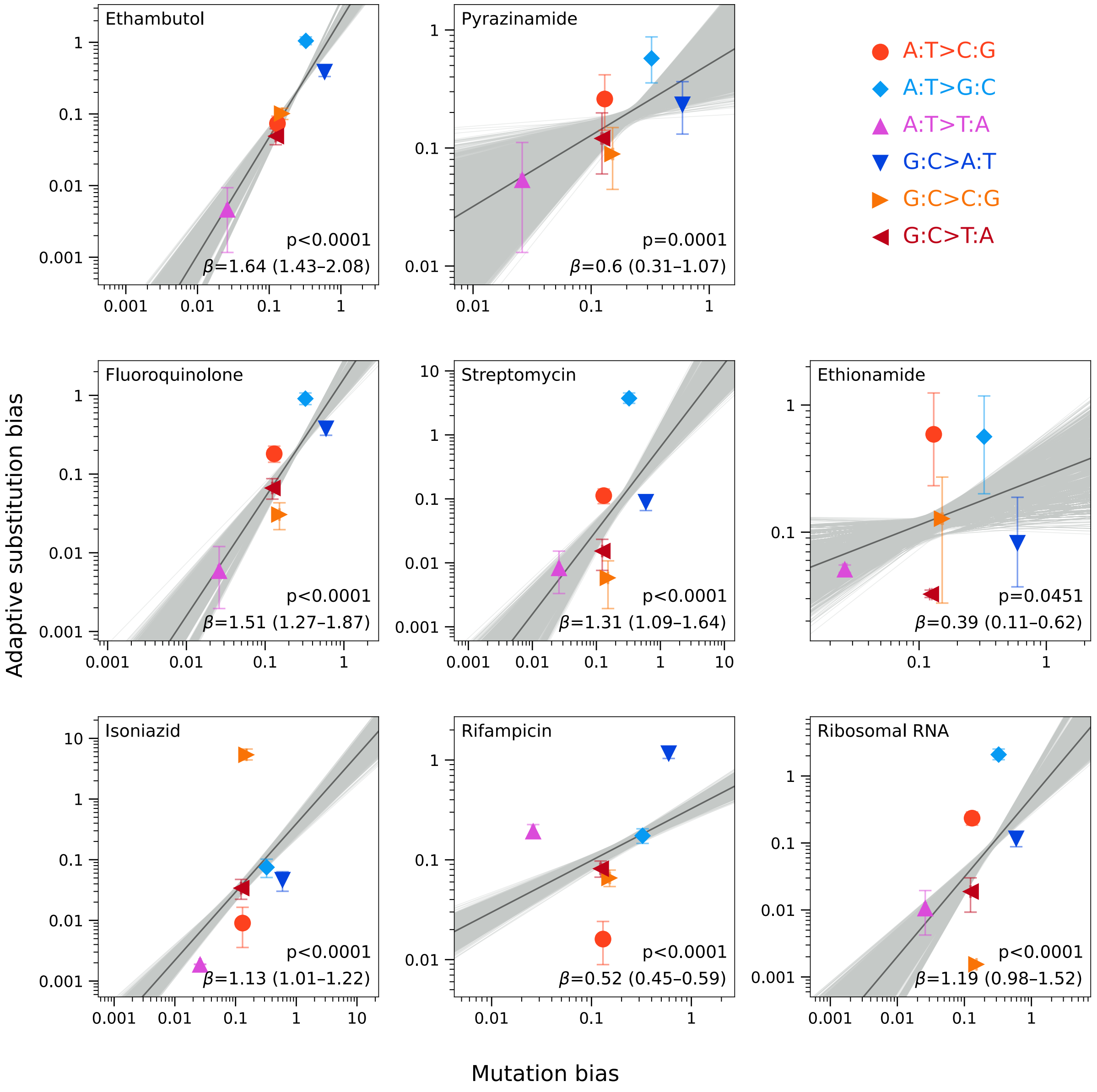


Fig. S12: **Bias in the spectrum of adaptive substitutions in *M. tuberculosis* reflects mutation bias, across multiple individual selection conditions and adaptive loci**. The *M. tuberculosis* adaptive substitutions dataset was separated into seven selection conditions characterized by drugs or drug classes, in addition to ribosomal RNA. Adaptive substitution bias in the case of ribosomal RNA was derived using Eq. 4, with path densities $p$ computed using the overall nucleotide composition of the ribosomal RNA gene *rrs* instead of codon frequencies. Least squares regression analyses were repeated on log-transformed biases, similar to the total intraspecific variation in bias shown in Fig. S1, with regression on empirical data (solid black line) and 10,000 bootstrap datasets (gray lines). The effect of mutation bias on bias in adaptive substitutions, $\beta$, is shown for each plot, with 95% bootstrap confidence interval shown in parentheses. Significance was assessed by simulating datasets (n=10,000) under the null model in which mutation bias has no effect on bias in adaptive substitutions.

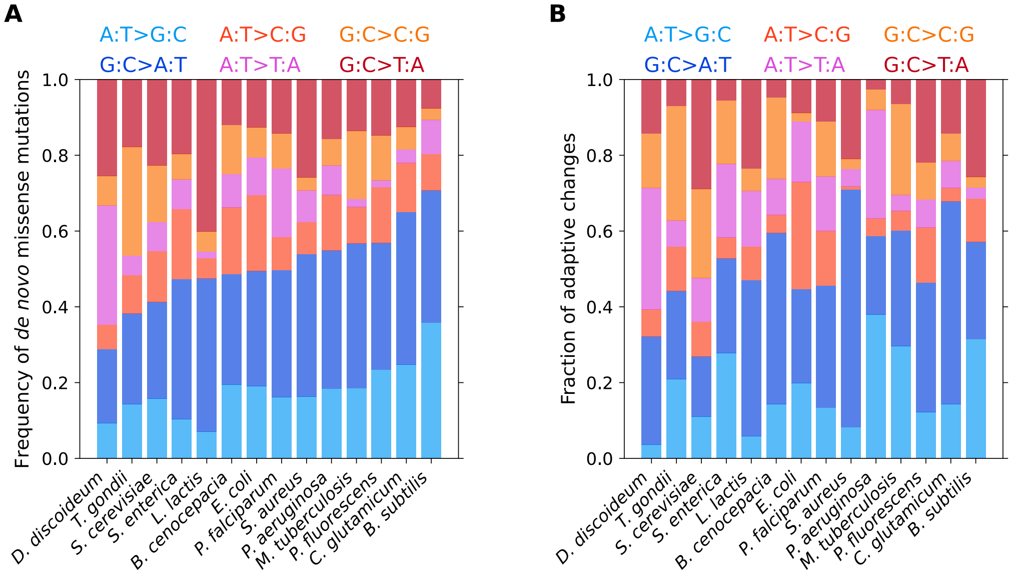


Fig. S13: **Mutation spectra and spectra of adaptive substitutions by species, in relative frequencies.** A) Frequency spectra of *de novo* missense mutations by species, arranged in order of increasing ratio of transitions (cool colors) to transversions (warm colors). To obtain missense mutation frequencies, mutation rates were weighted by the fraction of nonsynonymous mutational paths under that species' codon usage (Eq. 13). B) Frequency spectra of adaptive outcomes by species, arranged to preserve the ordering of species in panel A.

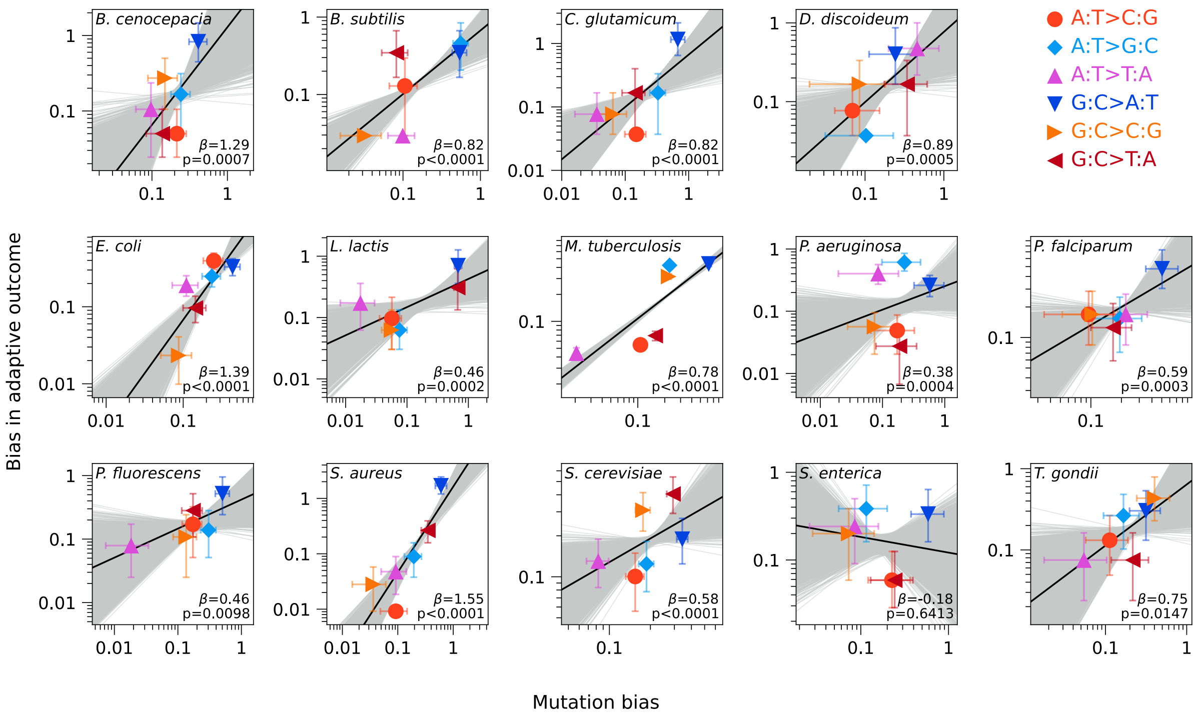


Fig. S14: **Biases in the relative frequencies of adaptive outcomes reflect biases in the relative frequencies of *de novo* mutations, in 13 of 14 species.** Bias in frequency among adaptive outcomes, for each of the six mutation classes, is plotted as a function of mutation bias. Biases are measured as frequency ratios of each of the mutation classes. A least squares regression on log-transformed empirical data yields a positive slope for 13 of 14 species. Regressions on empirical data are shown in solid black lines, and 10,000 bootstrap regressions are shown in gray lines. Significance was assessed by simulating datasets (n=10,000) under the null model in which mutation bias has no effect on bias in the frequencies of adaptive outcomes. See Table S6 for summary statistics.

| Species | $\beta$ | $\beta_{\text{null}}$ | Pearson's $r$ | p |
|---|---|---|---|---|
| *B. cenocepacia* | 1.29 (0.59–2.21) | -0.08 (-0.88–0.71) | 0.61 (0.27–0.83) | 0.0007 |
| *B. subtilis* | 0.82 (0.58–1.04) | -0.08 (-0.48–0.31) | 0.74 (0.51–0.87) | <0.0001 |
| *C. glutamicum* | 0.82 (0.46–1.24) | -0.07 (-0.52–0.39) | 0.74 (0.45–0.88) | <0.0001 |
| *D. discoideum* | 0.89 (0.38–1.43) | -0.14 (-0.68–0.46) | 0.73 (0.31–0.91) | 0.0005 |
| *E. coli* | 1.39 (1.08–1.82) | -0.16 (-0.41–0.08) | 0.79 (0.70–0.86) | <0.0001 |
| *L. lactis* | 0.46 (0.21–0.77) | -0.20 (-0.49–0.12) | 0.71 (0.33–0.91) | 0.0002 |
| *M. tuberculosis* | 0.78 (0.73–0.83) | 0.11 (0.07–0.14) | 0.81 (0.78–0.84) | <0.0001 |
| *P. aeruginosa* | 0.38 (0.08–0.73) | -0.13 (-0.42–0.16) | 0.22 (0.04–0.38) | 0.0004 |
| *P. falciparum* | 0.59 (0.19–1.04) | -0.35 (-0.87–0.15) | 0.76 (0.26–0.94) | 0.0003 |
| *P. fluorescens* | 0.46 (0.12–0.89) | -0.01 (-0.35–0.36) | 0.76 (0.19–0.94) | 0.0098 |
| *S. aureus* | 1.55 (1.29–1.93) | -0.20 (-0.46–0.07) | 0.86 (0.76–0.94) | <0.0001 |
| *S. cerevisiae* | 0.58 (0.22–0.97) | -0.48 (-0.84– -0.11) | 0.53 (0.20–0.76) | <0.0001 |
| *S. enterica* | -0.18 (-0.71–0.32) | -0.10 (-0.66–0.46) | -0.17 (-0.56–0.28) | 0.6413 |
| *T. gondii* | 0.75 (0.25–1.42) | 0.14 (-0.41–0.67) | 0.73 (0.24–0.92) | 0.0147 |

Table S6: **Summary statistics on intraspecific regressions for bias in frequencies of adaptive outcomes.** Results reflect the regression of biases in the adaptive substitution frequency spectrum on biases in the mutation frequency spectrum, for each of the 14 species (Fig. S14). Least squares regression analyses were performed on log-transformed data. Results include the statistic that captures the effect of mutation bias, $\beta$, on biases in adaptive outcomes, for the empirical data and null expectation in which mutation bias has no effect, the Pearson's correlation coefficient, followed by the p-value for the hypothesis that $\beta_{\text{empirical}} > \beta_{\text{null}}$. Columns for $\beta$ and Pearson's $r$ show estimates for the empirical data with 95% bootstrap confidence intervals in parentheses. The column for $\beta_{\text{null}}$ shows the point estimates for the null model in which codon frequencies determine adaptive substitution frequencies, with 95% confidence intervals from simulated datasets (n=10,000) under this null model shown in parentheses. Significance was assessed by comparison of the empirical point estimates to these null-model simulations in which mutation bias has no effect on bias in frequencies of adaptive outcome.

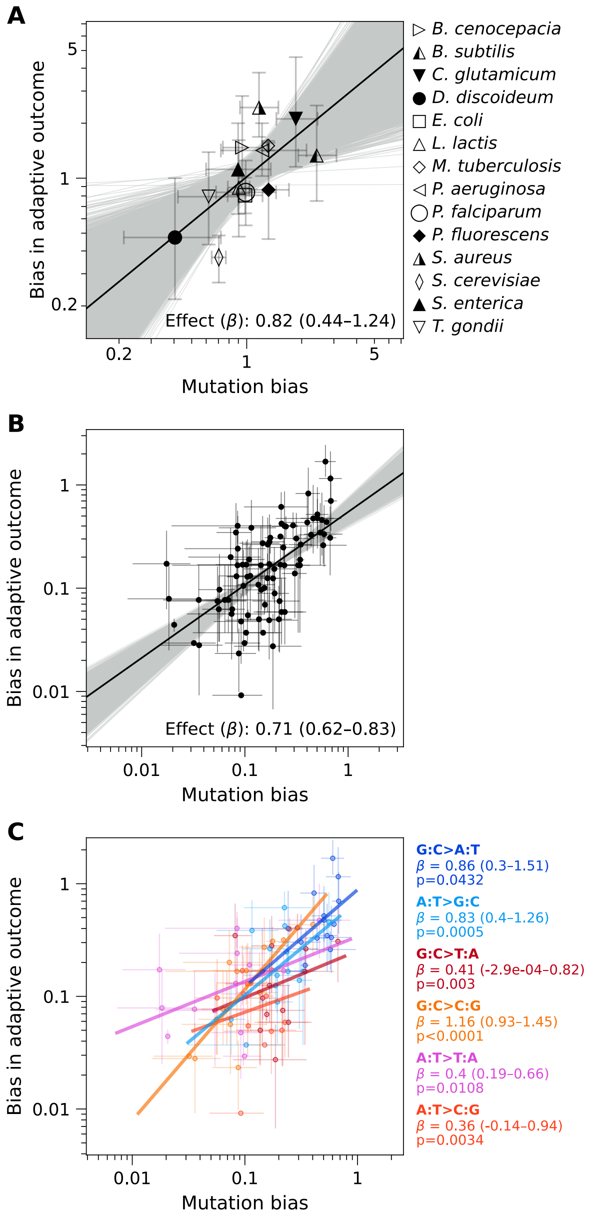


Fig. S15: **Bias in relative frequencies of adaptive outcomes reflects bias in relative frequencies of *de novo* mutations.** Regression analyses for transition bias (Fig. 3), overall bias across taxa and mutational classes (Fig. 4), and each mutation class individually (Fig. 5) are repeated in panels A, B, and C, respectively, using bias in frequencies rather than rates (see Fig. S13). Regressions are on log-transformed empirical data (solid black line for panels A and B; color-coded lines by mutation class for panel C), and on 10,000 bootstrap datasets (gray lines in panels A and B; not shown in panel C). The effect of mutation bias, $\beta$, is shown for each regression, with 95% bootstrap confidence interval in parentheses. Significance was assessed by simulating datasets (n=10,000) under the null model in which mutation bias has no effect on bias in adaptive substitutions (p=0.0004 for panel A, p<0.0001 for panel B, and individual mutation class-specific p-values shown in panel C).

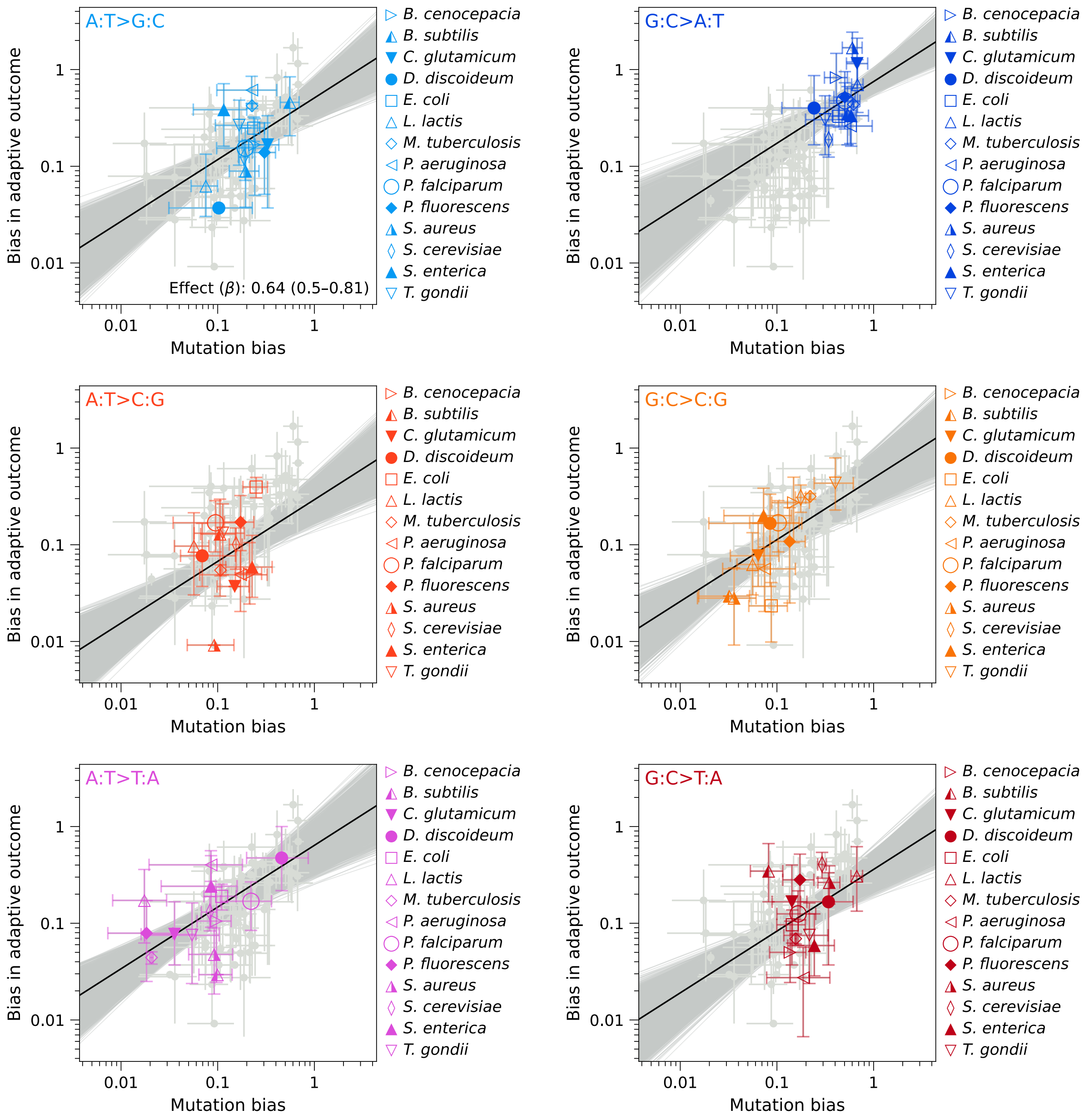


Fig. S16: **Variation across species in bias in adaptive frequency spectra reflects variation in bias in mutation spectra.** Bias in frequencies of adaptive outcomes plotted against mutation bias, across 14 species, for each of the six single-nucleotide mutation classes. The regression analysis shown here isolates interspecific variation using a single mutation bias effect parameter $\beta$ influenced by all mutation classes, and class-specific intercepts, on log-transformed empirical data (solid black lines) and 10,000 bootstrap datasets (gray lines), similar to Fig. S2 but with bias measured with respect to relative frequencies of adaptive outcomes and missense mutations. The effect parameter $\beta$ is shown in the plot for A:T>G:C (upper left) with the 95% bootstrap confidence interval in parentheses. Significance was assessed by simulating datasets (n=10,000) under the null model in which mutation bias has no effect (that is, biases in adaptive outcomes reflect biases in the fraction of accessible nonsynonymous paths), p<0.0001.

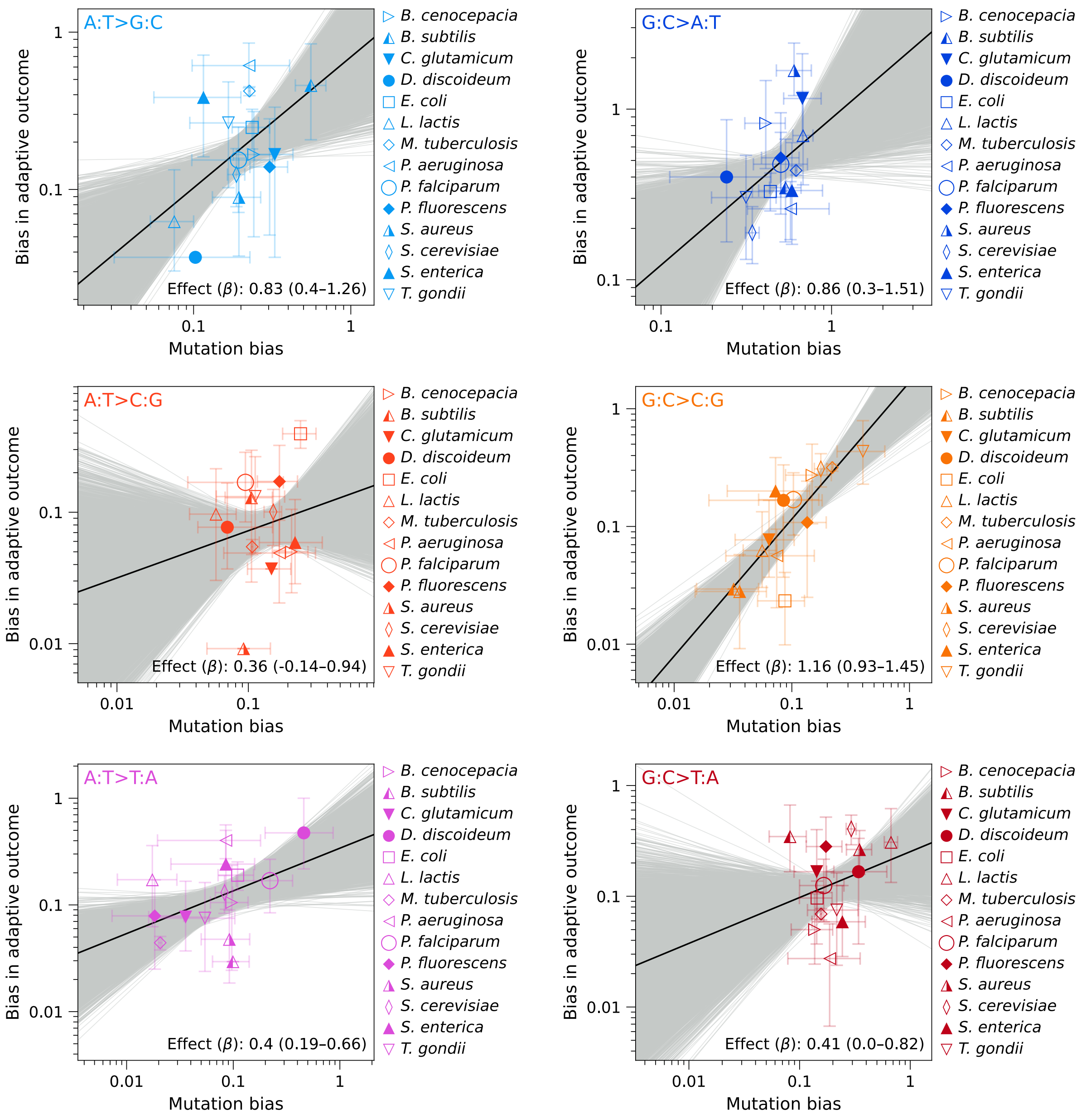


Fig. S17: **Variation across species in bias in frequencies of adaptive outcomes reflects variation in mutation bias, for each of the single-nucleotide mutation classes.** Biases in the frequencies of adaptive outcomes are plotted as a function of mutation bias, across 14 species. The data and regression analyses (solid black lines) are the same as in Fig. S15C, but separated by mutation class into individual plots, with bootstrap regressions (n=10,000) shown as gray lines. The 95% bootstrap confidence interval for mutation bias effect, $\beta$, is shown in parentheses for each regression. Significance (shown in Fig. S15C and Table S7) was assessed by simulating datasets (n=10,000) under the null model in which mutation bias has no effect (that is, biases in adaptive outcomes reflect biases in the fraction of accessible nonsynonymous paths).

| Mutation or bias type | Combined (intra- & interspecies) and mutation class-specific effects: $\beta$ | $\beta_{\text{null}}$ | $\beta_0$ | Pearson's $r$ | p | Isolation of interspecies variation in bias, aggregating across the six mutation classes: $\beta$ | $\beta_{\text{null}}$ | $\beta_0$ | p |
|---|---|---|---|---|---|---|---|---|---|
| Transition | 0.82 (0.44–1.24) | 0.0091 (-0.42–0.43) | 0.0077 (-0.05–0.07) | 0.69 (0.37–0.82) | 0.0004 | N/A | N/A | N/A | N/A |
| Intra- & inter-species | 0.71 (0.62–0.83) | -0.1 (-0.19–0.02) | -0.26 (-0.34– -0.19) | 0.61 (0.51–0.65) | <0.0001 | N/A | N/A | N/A | N/A |
| A:T>C:G | 0.36 (-0.14–0.94) | -0.38 (-0.76–0.14) | -0.78 (-1.26– -0.29) | 0.18 (-0.07–0.44) | 0.0034 | 0.64 (0.5–0.81) | 0.02 (-0.13–0.18) | -0.53 (-0.7– -0.39) | <0.0001 |
| A:T>G:C | 0.83 (0.4–1.26) | -0.12 (-0.58–0.4) | -0.16 (-0.49–0.13) | 0.53 (0.24–0.69) | 0.0005 | | | -0.29 (-0.43– -0.17) | |
| A:T>T:A | 0.4 (0.19–0.66) | 0.14 (-0.09–0.36) | -0.47 (-0.74– -0.2) | 0.45 (0.21–0.63) | 0.0108 | | | -0.19 (-0.39– -0.01) | |
| G:C>A:T | 0.86 (0.3–1.51) | 0.15 (-0.63–0.97) | -0.05 (-0.24–0.14) | 0.44 (0.15–0.65) | 0.0432 | | | -0.12 (-0.2– -0.04) | |
| G:C>C:G | 1.16 (0.93–1.45) | 0.17 (-0.1–0.46) | 0.23 (-0.04–0.48) | 0.82 (0.64–0.87) | <0.0001 | | | -0.31 (-0.49– -0.14) | |
| G:C>T:A | 0.41 (-0.00029–0.82) | -0.27 (-0.73–0.22) | -0.6 (-0.93– -0.33) | 0.26 (-0.00019–0.46) | 0.0030 | | | -0.44 (-0.6– -0.32) | |

Table S7: **Summary statistics on key results using bias in frequencies.** Summary statistics are shown for the regression analyses in Figs. S15 to S17. Regression analyses were performed on log-transformed data. Results include the statistic that captures the effect of mutation bias, $\beta$, on bias in frequencies of adaptive outcomes, for empirical data and null expectation in which mutation bias has no effect, the intercept parameter $\beta_0$, the Pearson's correlation coefficient, and the p-value for the hypothesis that $\beta_{\text{empirical}} > \beta_{\text{null}}$. Columns for $\beta$ and $\beta_0$, and Pearson's $r$ show estimates for the empirical data with 95% bootstrap confidence intervals in parentheses. The columns for $\beta_{\text{null}}$ show the point estimates for the null model in which codon frequencies determine adaptive substitution frequencies, with 95% confidence intervals from simulated datasets (n=10,000) under this null model shown in parentheses.

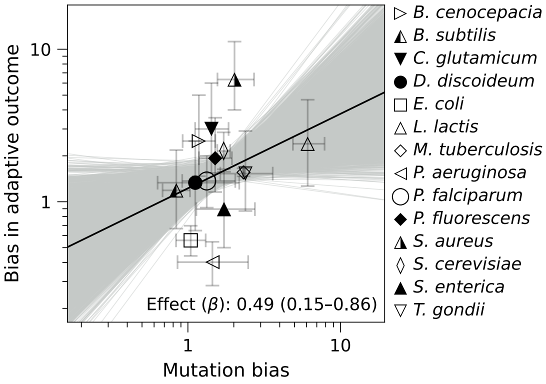


Fig. S18: **Mutational site bias contributes to site bias in frequencies of adaptive outcomes.** Site bias among adaptive outcomes is plotted against site bias in *de novo* missense mutations, using least squares regression on the log-transformed empirical data (solid black line) and 10,000 bootstrap datasets (gray lines). The 95% bootstrap confidence interval for mutation bias effect, $\beta$, is shown in parentheses. Significance was assessed by simulating datasets (n=10,000) under the null model in which mutation bias has no effect (that is, biases in adaptive outcomes reflect biases in the fraction of accessible nonsynonymous paths), p=0.0019.

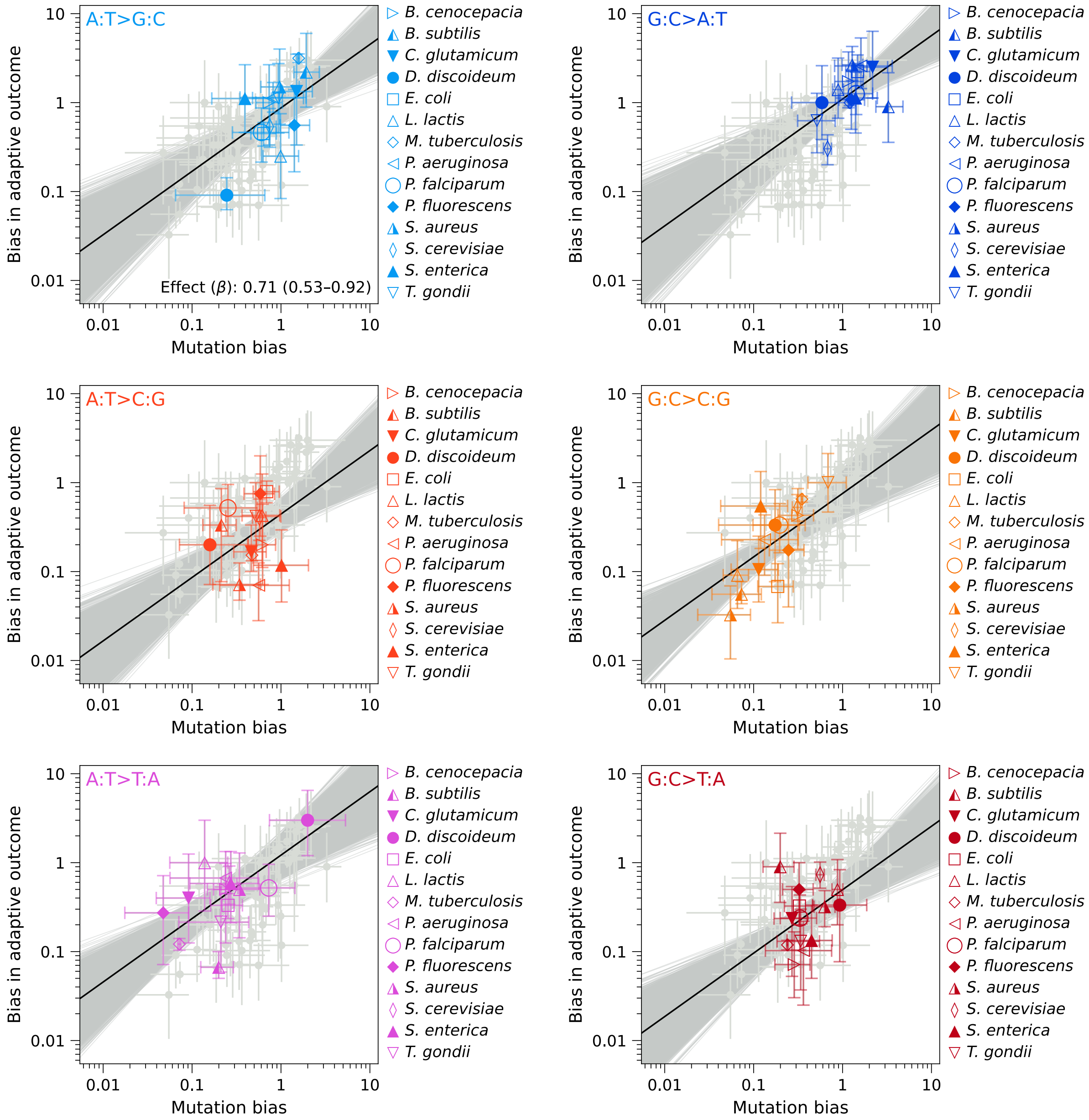


Fig. S19: **Variation in alternate-base bias contributes strongly toward the effect of mutation bias on frequencies of adaptive outcomes.** Bias in frequencies of adaptive outcomes within a species is plotted as a function of the corresponding mutation bias, across 14 species, for each of the six single-nucleotide mutation classes. This is similar to Fig. S16, but plotted values represent bias among the alternate (mutant allele) bases, conditioned on the identity of the mutated reference base, so that the magnitude of bias is unaffected by variation in underlying base mutability (Fig. S18, see Methods for details). Regression analysis was performed using a single mutation bias effect parameter, $\beta$, and class-specific intercepts, on log-transformed empirical data (solid black line) and 10,000 bootstrap datasets (gray lines). The effect parameter $\beta$ is shown in the plot for A:T>G:C (upper left) with the 95% bootstrap confidence interval in parentheses. Significance was assessed by simulating datasets (n=10,000) under the null model in which mutation bias has no effect (that is, biases in adaptive outcomes reflect biases in the fraction of accessible nonsynonymous paths), p<0.0001.

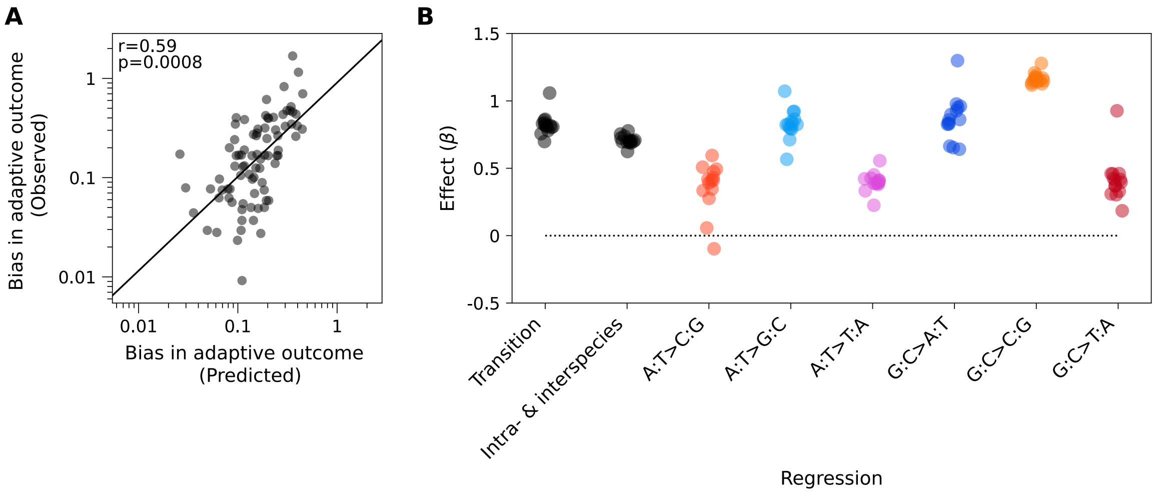


Fig. S20: **The relationship between mutation bias and adaptive substitution bias is robust under species exclusion when biases are expressed in terms of frequencies rather than rates.** A) Observed versus predicted bias in frequencies of adaptive substitution under leave-one-species-out cross-validation, aggregated across all 14 species and all six mutation classes. The approach is the same as in Fig. 6A but applied to frequencies instead of rates. The p-value for the leave-one-species-out correlation was assessed by permuting the species labels of the observed adaptive spectra and computing the Pearson's $r$ between the predicted and observed biases (n=10,000). B) Frequency-based leave-one-species-out estimates of $\beta$ were obtained by repeating the regression analyses reported in Fig. S15, while omitting each of the 14 species one at a time. See also Table S8.

| Held-out species | Transition | Intra- & interspecies | Interspecies | A:T>C:G | A:T>G:C | A:T>T:A | G:C>A:T | G:C>C:G | G:C>T:A |
|---|---|---|---|---|---|---|---|---|---|
| *B. cenocepacia* | 0.836 | 0.696 | 0.641 | 0.508 | 0.847 | 0.408 | 0.960 | 1.132 | 0.310 |
| *B. subtilis* | 1.058 | 0.693 | 0.691 | 0.407 | 0.796 | 0.451 | 0.899 | 1.153 | 0.926 |
| *C. glutamicum* | 0.756 | 0.695 | 0.648 | 0.410 | 0.920 | 0.390 | 0.664 | 1.172 | 0.459 |
| *D. discoideum* | 0.807 | 0.697 | 0.608 | 0.419 | 0.567 | 0.226 | 1.299 | 1.175 | 0.409 |
| *E. coli* | 0.817 | 0.687 | 0.595 | -0.097 | 0.823 | 0.387 | 0.837 | 1.141 | 0.396 |
| *L. lactis* | 0.818 | 0.778 | 0.718 | 0.595 | 0.712 | 0.556 | 0.828 | 1.170 | 0.185 |
| *M. tuberculosis* | 0.803 | 0.699 | 0.601 | 0.335 | 0.802 | 0.333 | 0.928 | 1.149 | 0.369 |
| *P. aeruginosa* | 0.809 | 0.724 | 0.627 | 0.431 | 0.790 | 0.379 | 0.977 | 1.147 | 0.370 |
| *P. falciparum* | 0.818 | 0.709 | 0.667 | 0.474 | 0.830 | 0.412 | 0.862 | 1.159 | 0.417 |
| *P. fluorescens* | 0.863 | 0.739 | 0.676 | 0.276 | 0.922 | 0.422 | 0.859 | 1.189 | 0.458 |
| *S. aureus* | 0.780 | 0.624 | 0.575 | 0.057 | 0.825 | 0.426 | 0.641 | 1.117 | 0.329 |
| *S. cerevisiae* | 0.698 | 0.708 | 0.599 | 0.346 | 0.825 | 0.400 | 0.654 | 1.127 | 0.303 |
| *S. enterica* | 0.832 | 0.754 | 0.690 | 0.494 | 1.072 | 0.388 | 0.951 | 1.208 | 0.460 |
| *T. gondii* | 0.855 | 0.704 | 0.613 | 0.391 | 0.864 | 0.391 | 0.827 | 1.278 | 0.424 |

Table S8: **Robustness of results to exclusion of taxa, for regressions based on frequencies rather than rates.** The values here are the ones plotted in Fig. S20B, along with (in column 4) the same iterative leave-one-out approach applied to the regression that isolates total interspecies variation on single-nucleotide mutation classes (Fig. S16), to assess the impact of omitting individual species on the estimated effect of mutation bias on frequencies of adaptive outcome.

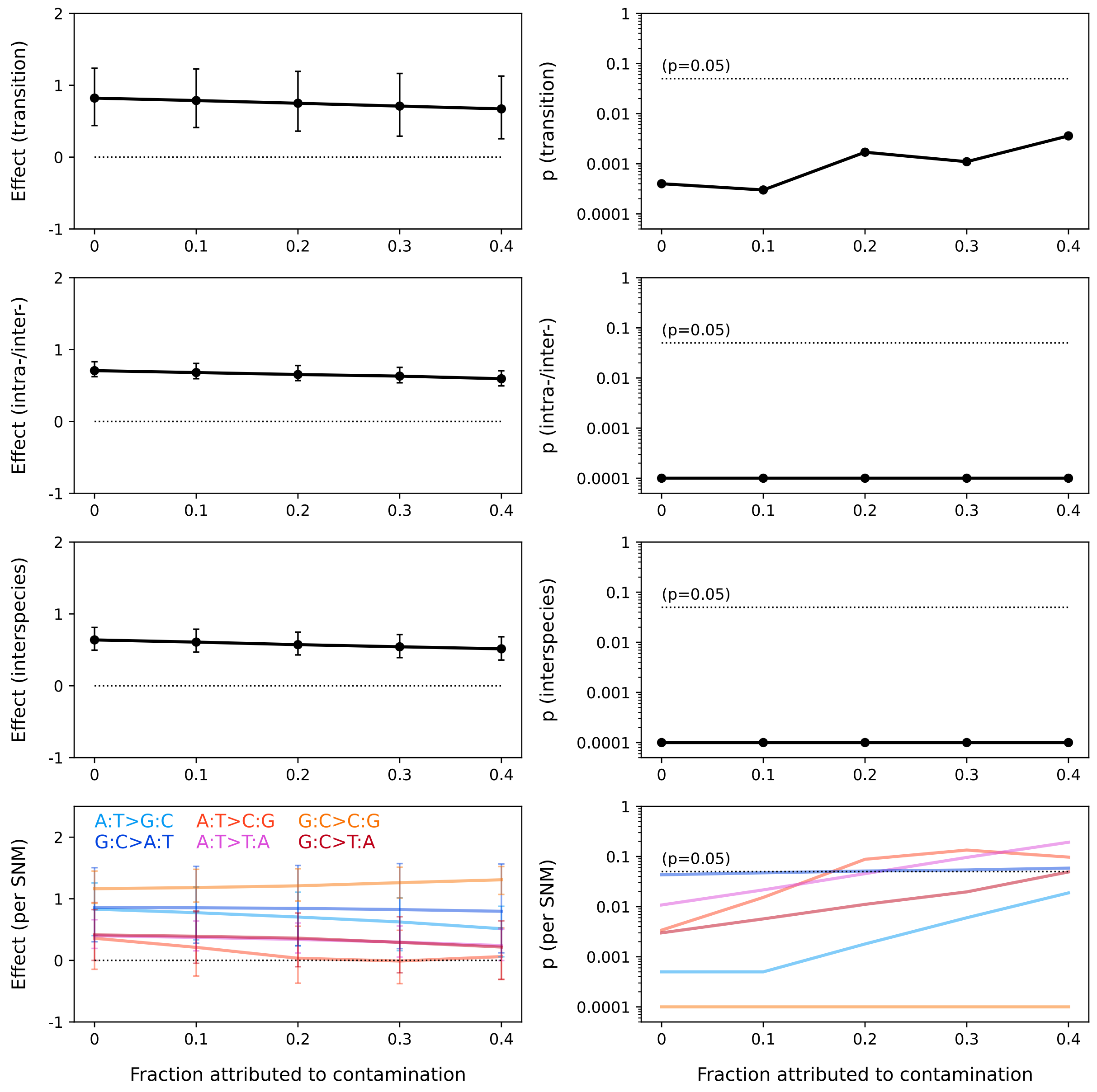


Fig. S21: **Robustness of results from regressions focusing on frequencies of adaptive outcome to contamination of the adaptive substitutions dataset.** Values plotted are the statistic characterizing the effect of mutation bias, $\beta$, on frequencies of adaptive outcome (left column) and the corresponding p-values (right column), upon increasing attribution of the adaptive substitutions dataset to non-adaptive contamination, under the assumption that the contamination spectrum (that is, the distribution of non-adaptive contaminants across the mutation classes) matches the *de novo* mutation spectrum. Regressions included in this analysis (by row, from top) are: transition bias, combined effect of intra- and interspecies variation in bias for all single-nucleotide mutation classes, the combined effect of interspecies variation across all six mutation classes, and the regressions on interspecies variation for each of the six mutation classes individually (Fig. S15A, S15B, S16, and S15C, respectively). Error bars show the 95% bootstrap confidence interval. Significance was assessed by simulating datasets (n=10,000) under the null model in which mutation bias has no effect (that is, biases in adaptive outcomes reflect biases in the fraction of accessible nonsynonymous paths).
